# Development in context: interferon response networks regulate human fetal thymic epithelial cell differentiation

**DOI:** 10.1101/2022.10.02.510339

**Authors:** Abdulvasey Mohammed, Benjamin D. Solomon, Priscila F. Slepicka, Kelsea M. Hubka, Hanh Dan Nguyen, Wenqing Wang, Martin Arreola, Michael G. Chavez, Christine Y. Yeh, Doo Kyung Kim, Virginia Winn, Casey A. Gifford, Veronika Kedlian, Jong-Eun Park, Georg A. Hollander, Vittorio Sebastiano, Purvesh Khatri, Sarah A. Teichmann, Andrew J. Gentles, Katja G. Weinacht

**Affiliations:** Department of Pediatrics, Division of Stem Cell Transplantation and Regenerative Medicine, Stanford School of Medicine, Stanford CA, USA; Department of Pediatrics, Division of Allergy and Immunology, Stanford School of Medicine, Stanford CA, USA; Department of Bioengineering, Stanford University, Stanford, CA, USA; Department of Biomedical Data Science, Stanford University, Stanford, CA, USA; Department of Obstetrics and Gynecology, Stanford School of Medicine, Stanford CA, USA; Department of Pediatrics, Division of Cardiology, Stanford School of Medicine, Stanford CA, USA; Wellcome Sanger Institute, Cambridge, UK; Department of Pediatrics and Institute of Developmental and Regenerative Medicine, University of Oxford, Oxford, United Kingdom; Institute for Immunity, Transplantation, and Infection, Stanford University, Stanford, CA, USA; Department of Pathology, Stanford School of Medicine, Stanford, CA, USA

## Abstract

The thymus instructs T cell immunity and central tolerance, yet its therapeutic potential remains untapped. The quest to regenerate thymic function for clinical application is lagging while the signals that drive thymic epithelial cell differentiation remain incompletely understood. Here, we elucidate pathways instructing commitment and specialization of the human thymic epithelial stroma through complementary single cell transcriptomic approaches. First, we identify gene regulatory networks that define fetal thymic epithelium in the thus far unexplored context of other anterior foregut-derived organs; then, we characterize lineage trajectories within the thymic epithelial compartment across embryonic, fetal, and early postnatal stages. Activation of interferon response gene regulatory networks distinguished epithelial cells of the thymus from those of all other anterior foregut-derived organs. Interferon signals were processed differentially within thymic cortical and medullary lineages, reflected in distinct *NFκB* and *IRF* signatures, respectively. Our study reveals novel, translatable insights into the developmental programs underlying thymic epithelial cell differentiation that may advance the field of regenerative cell therapies.

**SUMMARY:** Single cell transcriptomics of anterior foregut-derived organs identifies pathways governing thymic epithelial commitment and specialization.

## INTRODUCTION

The thymic epithelium comprises a highly specialized set of cells that attract lymphoid progenitors, promote their proliferation and maturation into thymocytes, and facilitate the selection of a diverse, self-tolerant T cell receptor repertoire^1–4^. As such, the thymus is central to the development of the adaptive immune system. Thymic function naturally declines in the second decade of life^5^. In addition, thymic output is susceptible to a wide range of pathogenic insults including medications, radiation, infection and graft-versus-host disease^2,3,6,7^. Endogenous or acquired thymic compromise leads to immunodeficiency, autoimmunity and inflammation^7–9^. Regenerating thymic function, such as through induced pluripotent stem cell (iPSC)-derived thymic epithelial cells, holds great therapeutic promise.

The thymus is derived from the anterior foregut endoderm. During development, the anterior foregut gives rise to the ventrally located respiratory system and the dorsally located esophagus^10^. The anterior-most segment of the foregut develops into the pharyngeal apparatus. Essential to the development of the mammalian pharyngeal apparatus is the out-pocketing of the pharyngeal pouches^11^. The thymic primordium, also referred to as thymus anlage, forms from the ventro-caudal region of the third pharyngeal pouch, while the dorsal aspect gives rise to the parathyroid (Fig. 1A). The signals that induce iPSCs into anterior foregut endoderm have long been described and protocols to differentiate anterior foregut endoderm further into epithelial cells of lung and esophagus *in vitro* have successfully been established^12–17^. Recent studies of murine and human thymus using single cell transcriptomics have greatly expanded our understanding of thymic epithelial cell (TEC) diversity^18–27^, yet the signals that drive TEC fate beyond the anterior foregut stage remain incompletely understood. Elucidating these signals is essential to derive TECs from iPSCs in an effort to enable regenerative cell therapies.

**Figure 1.**
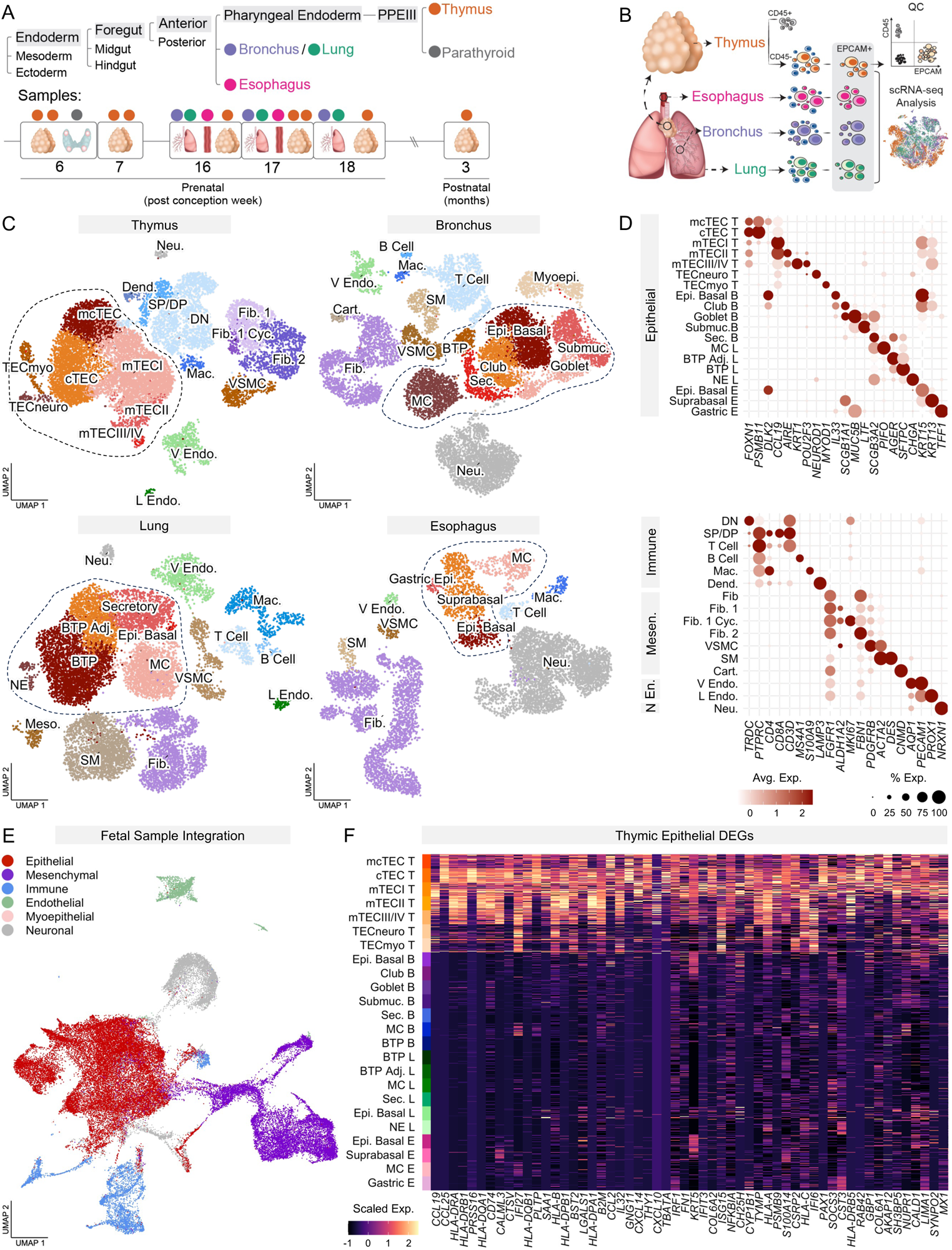
Cellular composition of human fetal anterior foregut-derived tissues. (A) Ontogeny of anterior foregut-derived organs. Thymus, bronchus, lung, esophagus and parathyroid arise from the endodermal germ layer (PPEIII, third pharyngeal pouch endoderm) (top). Summary of samples including developmental age and tissue of origin. Colors denote tissue type (thymus, orange; bronchus, blue; lung, green; esophagus, pink; parathyroid, grey) (bottom). (B) Workflow of cell processing and quality control prior to transcriptional profiling. (C) UMAP visualization of human anterior foregut-derived tissues colored by cell type (DN, double negative thymocytes; SP/DP, double positive/single positive thymocytes; Mac, macrophages; Dend, dendritic cells; Fib, fibroblasts; Cyc, cycling; VSMC, vascular smooth muscle cells; SM, smooth muscle cells; Cart, cartilage; V Endo, vascular endothelial cells; L Endo cells, lymphatic endothelial cells; Neu, neuronal cells; Epi, epithelial cells; Submuc, submucosal cells; Sec, secretory cells; MC, multiciliated; BTP, bud tip progenitor cells; adj, adjacent; NE, neuroendocrine cells). Dotted line denotes epithelial clusters. (D) Dot plot of select, cell type-defining genes in individual anterior foregut-derived tissues. Here and onwards, color represents average expression of marker genes in each cell type. Dot size represents percentage of cells in each cell type expressing the selected marker gene (T, thymus; B, bronchus; L, lung; E, esophagus; Mesen, mesenchymal; En, endothelial; N, neuronal). (E) UMAP visualization of integrated anterior foregut-derived tissues at the fetal stage of development (horizontal integration), colored by organ of origin. (F) Heat map of the top thymic epithelial differentially expressed genes (DEGs) across all epithelial clusters from (C) downsampled to 30 cells per cluster.

In human embryonic samples, the third pharyngeal pouch transcriptional regulators HOXA3, PAX9, TBX1 and FOXG1 are detected as early as the beginning of post conception week (pcw) 6 and precede the expression of the master thymic transcription factor FOXN1 mid-week 6^28^. Lymphoid progenitors enter the thymus anlage as early as pcw 7^21,28^. At pcw 8, less than 5% of cells in the developing human thymus are of hematopoietic origin (CD45^+^)^23^. In contrast, by pcw 10, more than 90% of cells in the thymus are of hematopoietic orgin^23^, highlighting the dramatic expansion of thymocytes (i.e. immature T cell precursors) and other immune cells during thymic development.

Important insights into the developmental programs that instruct thymic organogenesis have been derived from mice with biallelic loss of function mutations in Foxn1, which result in an athymic and hairless (nude) phenotype^29,30^. Thymic development in nude mice arrests before colonization with lymphoid progenitors occurs^31,32^. The thymic rudiment of Foxn1^null^ mice displays nonfunctional cystic features strongly resembling respiratory epithelium in morphology and gene expression. It has therefore been suggested that Foxn1 overwrites respiratory cell fate as the default pathway in the anterior foregut^32–35^. Notably, NOD-scid IL2Rg^null^ (NSG) mice with genetically impaired immune functions of hematopoietic origin leading to a lack of thymocytes also display distorted thymic architecture with cystic features resembling respiratory epithelium^36^. Although NSG mice do not have cell-intrinsic genetic defects in thymic epithelial cell differentiation, they often exhibit permanently impaired thymic function leading to poor T cell reconstitution even if healthy hematopoietic stem cells are transplanted later in life^36,37^. This suggests that Foxn1 is necessary but not sufficient to fully induce thymic epithelial cell fate. While thymic defects in NSG mice^36^ support that development of thymic epithelial and lymphoid niches during embryogenesis are intertwined, the corresponding gene regulatory networks governing the lymphoid-driven commitment and maturation of the thymic epithelial stroma have not yet been characterized.

Here we identify regulators underlying thymic epithelial differentiation by comparing epithelial cells from human fetal organs of shared anterior foregut origin (thymus, bronchus, lung, esophagus and parathyroid) using single cell transcriptomics. We characterize developmental programs unique to specific TEC subsets by analyzing human samples from embryonic, fetal, and early postnatal stages. Our studies yield translatable insights designed to advance regenerative cell therapy approaches.

## RESULTS

### Human fetal anterior foregut-derived organs at single cell resolution

Single cell RNA-sequencing (scRNA-seq) was performed on 17 samples from 15 distinct biological specimens. For the comparison of human fetal anterior foregut-derived organs, we sampled midgestation tissues from thymus (pcw 16, 17, 18), airway/bronchus (pcw 16, 17, 18), lung parenchyma (pcw 16, 17,18) and esophagus (pcw 16 and 17). We refer to the comparison of different tissues from the same stage of development as *horizontal* comparison. We also performed *vertical* comparisons between human thymus samples obtained during early gestation (pcw 6, 7), midgestation (pcw 16, 17, 18) and postnatally (age 3 months) to examine lineage trajectories and epithelial cell diversification within the thymus (Fig. 1A). The earliest thymus samples were used unmanipulated due to very low numbers of thymocytes at this stage of development. At later stages, thymic samples were depleted of CD45^+^ thymocytes. All samples were then enriched for EPCAM^+^ epithelial cells (Fig. 1B). Sample characteristics are summarized in Supplementary Table S1.

Preprocessing and integration of scRNA-seq samples using Seurat yielded a dataset of 52,162 single cell transcriptomes with a mean of 3522 unique genes per cell, and an average of 14,839 total counts per cell. Cells within unsupervised clusters were annotated in concordance with previously published human single cell datasets for thymus^21^, respiratory system^38,39^ and esophagus^40^, respectively.

In fetal thymus, we identified 18 epithelial and non-epithelial cell populations (Fig. 1C). Epithelial cells included cortical thymic epithelial cells (cTEC) and medullary thymic epithelial cells (mTECI). An intertypic mcTEC^21^ population with dim expression of cTEC and mTEC markers and uniquely high expression of *CTGF* and *DLK2,* was the third most abundant population. mTECII were characterized by high expression of *AIRE* and *CLDN4*. The medullary TEC subpopulations mTECIII and mTECIV were defined by expression of *KRT1* and *POU2F3,* respectively, which correspond to the post-*AIRE* keratinocyte-like mTEC and Tuft cells in the mouse^19,20,41,42^. The recently characterized neuroendocrine (TECneuro) and muscle-like (TECmyo) TEC populations were also observed^21^. Immune cell clusters within the thymus included a spectrum of developing thymocytes from double-negative (DN) through double-positive (DP) thymocytes to single-positive stages, thymus-resident macrophages and dendritic cells. Other nonepithelial clusters included vascular smooth muscle cells, aldehyde dehydrogenase-positive fibroblasts (thymic fibroblasts type 1^21^), fibronectin-high fibroblasts (thymic fibroblasts type 2^21^), vascular endothelial cells, lymphatic endothelial cells and neuronal cells (Fig. 1C, D and fig. S1).

Tissues of the bronchus clustered into airway epithelial basal cells and epithelial cells from the adjacent respiratory parenchyma. In addition, the airway microenvironment was comprised of immune cells (B cells, T cells), myoepithelial cells, smooth muscle, vascular smooth muscle, fibroblasts, cartilage, endothelial cells and neuronal cells. Lung tissue expectedly shared many of the same cell types surrounding the airway, albeit in reverse ratios. Cell types not derived from the foregut included neuroendocrine cells, immune cells (B cells, T cells, macrophages), fibroblasts, mesothelial cells, vascular smooth muscle cells, smooth muscle cells, endothelial cells, lymphatic endothelial cells and neuronal cells. The esophageal tissue included epithelial basal cells, suprabasal epithelial and gastric epithelial cells. Nonepithelial cells of the esophagus were comprised of immune cells (T cells, macrophages), smooth muscle, vascular smooth muscle, fibroblasts, endothelial cells and a cluster of neuronal cells (Fig. 1C, D and fig. S1). All identified cell pollutions and their defining markers are summarized in Supplementary Table S2.

Integrated data sets from thymus, bronchus, lung and esophagus revealed shared epithelial, mesenchymal, immune, endothelial, myoepithelial and neuronal clusters regardless of organ of origin (Fig. 1E and fig. S2). To investigate human thymic epithelial fate in the thus far unexplored context of developmentally related epithelia, we subclustered epithelial cells from all anterior foregut-derived organs and identified top differentially expressed genes unique to the thymus. Expectedly, we detected key markers of thymic development and function, including *PAX1*, *PRSS16, CCL19, CCL25, TBATA*. Unexpectedly, we discovered numerous interferon-inducible or interferon-modulating genes including *IRF1, IFI6, IFI27, IFIT3, MX1, ISG15* and *NFKBIA* with uniquely high expression in epithelial cells of the thymus, raising the possibility that interferon (IFN) exerts unique effects in the developing thymus.

### Selective IFN response network activation in TECs

IFNs induce cell intrinsic programs mediating a wide range of biological responses, e.g. through the JAK-STAT pathway^43–45^ and NF-κB^46^. To explore the role of IFN signaling during thymic development, we investigated gene regulatory networks across anterior foregut-derived epithelial cells using the single cell regulatory network inference and clustering algorithm SCENIC^47, 48^. Modules, composed of a transcription factor and its inferred targets, are in the following referred to as regulons. UMAP visualization of organ-specific regulon expression distinctly separated the thymic epithelium from epithelial cells of bronchus, lung and esophagus (Fig. 2A). Across all epithelial cells, top differentially expressed regulons specific to the thymus encompassed a broad selection of inflammatory response regulators including IFN regulatory factors (*IRF1, IRF4, IRF6, IRF7, IRF9)*, signal transducer and activator of transcription factors (*STAT1, STAT2, STAT3, STAT5A*), nuclear factor-κB (NF-κB) complex components or modifiers thereof (*NFKB1, NFKB2, RELB, NFATC2, HIVEP1*^49^), and additional modulators of inflammatory response signaling such as *BHLHE40*^50^ and *BATF*^51^. Also identified were the key regulators of thymic patterning and development, *FOXN1*, *PAX9*, *FOXG1*^28^ and *TP63*^52^ (Fig. 2B).

**Figure 2.**
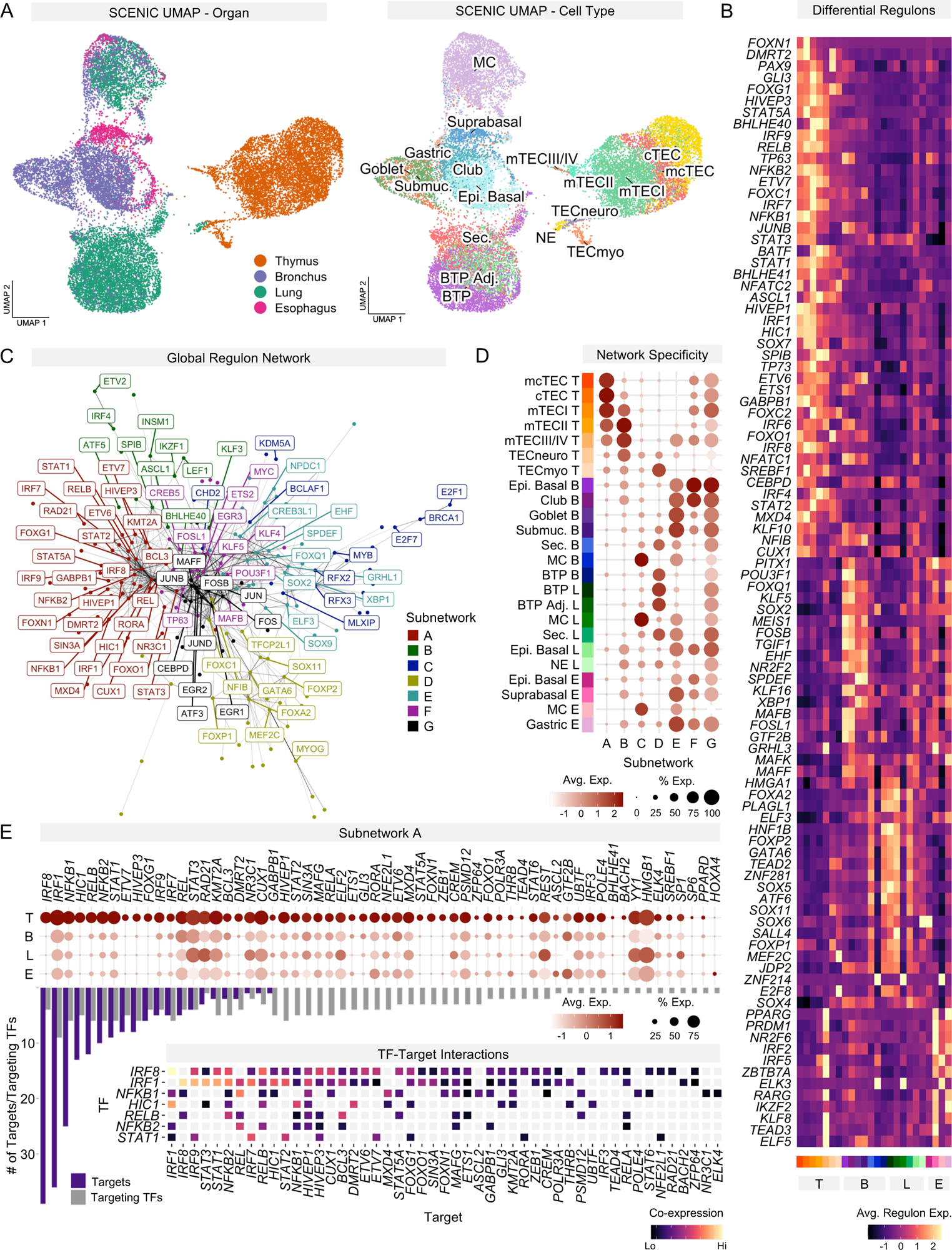
Differential gene regulatory analysis across anterior foregut-derived epithelial cells. **(A)** UMAP visualization of SCENIC transcription factor regulons in all fetal epithelial cells colored by tissue of origin (left) and cell type (right). **(B)** Heatmap of differentially expressed transcription factor regulons across anterior foregut-derived tissues (T, thymus; B, bronchus; L, lung; E, esophagus). Color key corresponds to annotated clusters from (D). **(C)** Visualization of transcription factor regulon subnetworks from (A) defined by Leiden algorithm. **(D)** Dot plot representing the average expression of individual transcription factor subnetworks in select epithelial cell subpopulations. **(E)** Dot plot of all subnetwork A genes across anterior foregut-derived tissues. Bar plot corresponds to number of target genes an individual transcription factor has (blue), and the number of regulons that target the respective transcription factor (gray) (top). Heatmap of top inferred interactions found in subnetwork A (TF, transcription factor) (bottom).

To investigate interactions between individual regulons, a global network of significant connections was constructed using the Leiden algorithm^53,54,55^. This process delineated 7 unique subnetworks (A-G) for which linkage is inferred (Fig. 2C and fig. S3). Modular expression of the transcription factors defining these subnetworks demonstrated organ and cell type specificity for select subnetworks (A-C) while others were broadly shared between organ systems (D-G) (Fig. 2D). Subnetworks A and B were unique to the thymus with subnetwork A defining the primary TEC populations cTEC, mcTEC and mTECI while subnetwork B defined mTECII, mTECIII/IV and TECneuro. Subnetwork A linked the IFN regulatory factor regulons *IRF1, IRF3, IRF7, IRF8* and *IRF9* with the STAT family regulons *STAT1, STAT2, STAT3 STAT5A,* and *STAT6,* the NF-κB family regulons *NFKB1, NFKB2, REL, RELA, RELB,* and the NF-κB modulators *HIVEP1*^49^ and *BHLHE40*^50^, all of which either transmit or modulate the cellular response to IFNs (Fig. 2E). *IRF8, IRF1* and *NFKB1* regulons comprised most of the interactions found in subnetwork A through co-expression with numerous additional *IRF*-, *STAT*-, and NF-κB-associated regulons unique to thymic epithelium. Notably, the developmental thymic transcription factors *FOXN1* and *FOXG1* were also part of the IFN response-centric subnetwork A and inferred targets of *IRF1* and *IRF8* (Fig. 2E). A biological association between *FOXG1* and IFN signaling has recently been made by a report describing that congenital viral infections recapitulate the spectrum of birth defects found in congenital *FOXG1* syndrome^56^. Together, these observations suggest that an intricate network of IFN regulated signaling pathways is selectively activated during TEC development but not in epithelial cells of other anterior foregut-derived organs.

### IFN responses in TECs increase with developmental age

To investigate if alternate methods of analysis would also demonstrate evidence of increased IFN response signaling in the thymic epithelium, we performed Gene Set Enrichment Analysis (GSEA) on select anterior foregut-derived epithelial clusters: mcTEC, cTEC, mTECI, mTECII, bronchial epithelial basal cells, esophageal epithelial basal cells and lung bud tip progenitor cells (Fig. 3A). GSEA network visualization of positively enriched gene sets revealed three main clusters (Fig. 3B and fig. S4, S5A). Within the cTEC/mTECI/mTECII cluster, mTECI was highly enriched for genes involved in IFN signaling, cytokine immune signaling, and antigen presentation. Gene sets with core enrichment of MHC genes constituted the remainder of gene sets within the cluster and were enriched in both cTEC and mTECII (Fig. S5B). In a separate cTEC-specific cluster, genes involved in TCR signaling, proteasome peptide processing, cross presentation of exogenous antigens, and IL-1 signaling were enriched. Enrichment of ribosomal and translational gene sets was shared between mcTEC, esophageal epithelial basal and lung bud tip progenitors. To gain a more granular understanding of the effects induced by IFNs we interrogated enriched IFN signaling gene sets for the most differentially expressed genes (Fig. 3C). In addition to the previously noted regulators and transducers of IFN signaling, cTEC, mTECI and mTECII exhibited differential expression of MHC class I and II genes, as well as the IFNγ receptor genes *IFNGR1* and *IFNGR2*. Notably, MHC class I and II are upregulated by IFNγ^57^.

**Figure 3.**
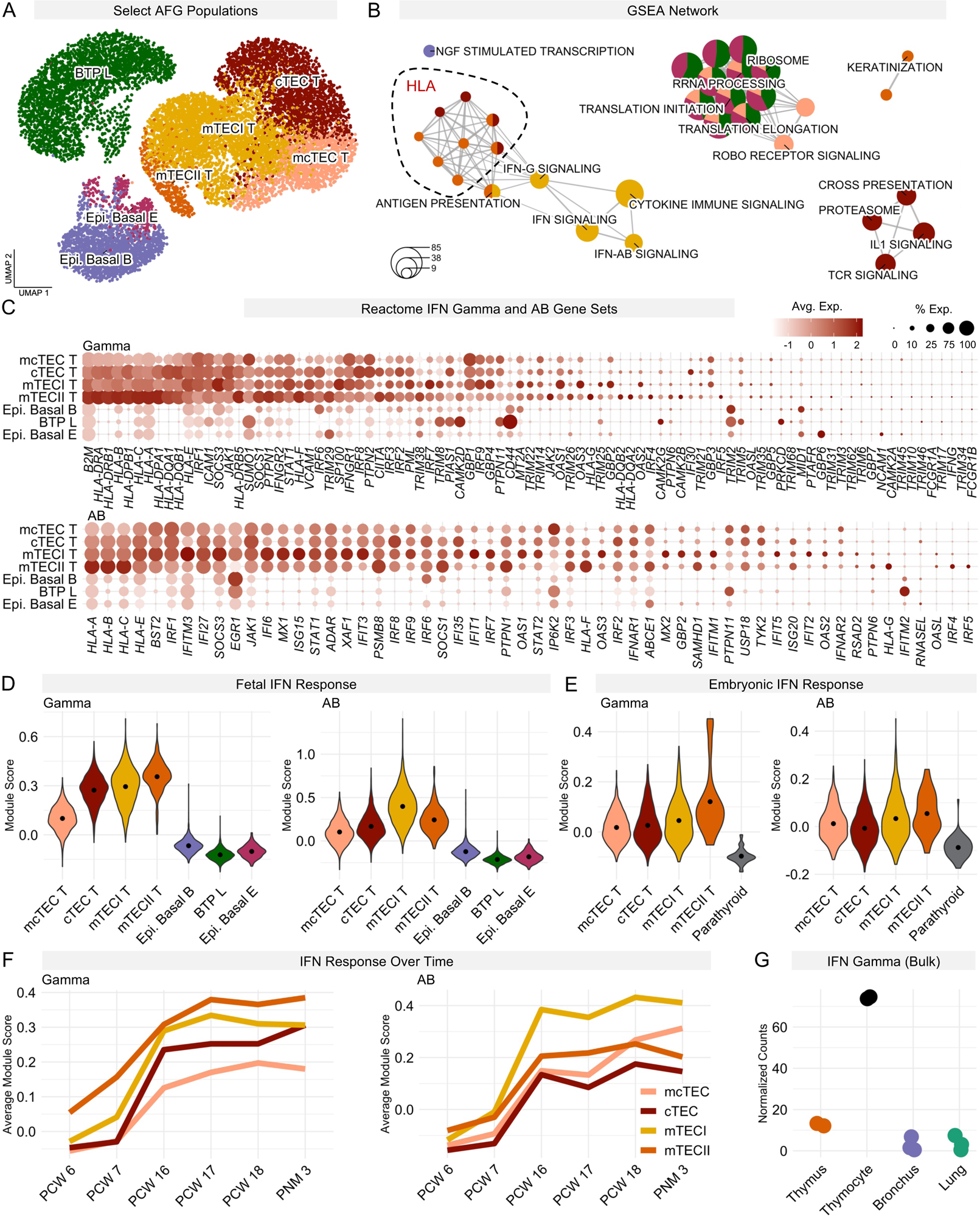
IFN response signatures in select epithelial cell subpopulations throughout development. **(A)** UMAP visualization of select epithelial cell subpopulations for thymus, bronchus, lung and esophagus at fetal stages of development. AFG, anterior foregut-derived epithelial cell. **(B)** Network visualization of differentially expressed GSEA gene sets across cell types from (A), colored accordingly. Each node represents a specific GSEA gene set (*p*<0.05). Edges between nodes denote a significant number of shared genes between nodes. Node size reflects the number of genes within each gene set. Gene set names have been abbreviated. Gene sets enriched primarily of *HLA* genes are within the dotted circle. **(C)** Dot plot of all detected Reactome-curated type I and II IFN response genes across populations from (A). **(D)** Violin plots depicting the average expression of Reactome-curated type I and II IFN response gene sets (modules) across populations from (A). Here and onwards, the dot within the violin plot indicates mean expression values. **(E)** Violin plot depicting the average expression of Reactome-curated type I and II IFN response modules in thymic and parathyroid epithelial cells at embryonic stages of development. **(F)** Average expression of Reactome-curated type I and II IFN response modules across embryonic, fetal and postnatal time points in select TEC subpopulations. **(G)** Cumulative (bulk) RNAseq expression of *IFNG* in thymocytes and EPCAM positive-selected epithelial cells from thymus, bronchus and lung.

IFNα and IFNβ are type I IFNs, IFNγ represents type II IFN^43,58^. To establish a quantifiable measure of IFN responses across queried cell types, we integrated IFN response gene sets into type I and II IFN specific expression modules, hereby referred to as module I and II. At midgestation (pcw 16-18), module I showed highest expression in mTECI while mTECII showed highest expression of module II. In contrast age-matched epithelial cell populations from bronchus, lung and esophagus showed significantly reduced expression across both modules (Fig. 3D). Early in gestation (pcw 6-7), mcTEC, cTEC, mTECI and mTECII showed higher expression of modules I and II relative to age-matched parathyroid epithelial cells (PTEC) (Fig. 3E). When compared longitudinally across samples of advancing gestational ages, modules I and II demonstrated an increasing trajectory over time (Fig. 3F) paralleling the age-dependent increase in the number of immune cells in the developing thymus.

Antigen presenting cells such as dendritic cells and macrophages are widely recognized producers of type I IFN^59^. Activated T cells, together with NK cells, are widely recognized as producers of type II IFN^60^, yet less is known about the capacity of developing T cells to produce type II IFN during human thymopoiesis. To investigate a possible link between the expansion of the lymphoid niche and the increase in type II IFN response in the thymic epithelium over time, we sought to quantify the relative contributions of immune and stromal cells to the production of *IFNG*. Because *IFNG* as well as *IFNA1, IFNA2* and *IFNB1* single cell transcripts largely evaded detection across our data set, we performed bulk RNA-sequencing on magnetic bead-enriched fractions of thymocytes, TEC, bronchial epithelial cells and lung epithelial cells. Thymocyte samples expressed notably higher levels of *IFNG* than epithelial cells from thymus, bronchus or lung suggesting thymocytes were primary producers of IFNγ in the thymus (Fig. 3G). The increase in IFN response during development combined with the higher expression of *IFNG* in thymocytes points to IFNγ as a mediator of the lymphoid-driven maturation of the thymic epithelium.

### Modifiers of IFN responses in thymic epithelial subsets

To examine TEC diversification throughout development we integrated TECs from samples of subsequent gestational ages (pcw 6, 7, 16, 17, 18) and 3 months post birth (*vertical* integration). UMAP visualization revealed a new TEC cluster in spatial proximity to mcTEC (Fig. 4A and fig. S6), not identified in our earlier analysis (*Figure 1*). The newly identified population clustered independently due to high differential expression of genes implicated in cell cycle, DNA synthesis and proliferation (*TOP2A, CENPF, NUSAP1, TUBAB1, MKI67*) (fig. S7 and S8) and demonstrated the highest average expression of G2M and S phase gene sets among all clusters (fig. S9). Accordingly, we designated this population cycling TEC (cyTEC). CyTEC were most predominant in the earliest (embryonic) samples and decreased in frequency with increasing developmental age. mTECII showed a relative increase in frequency over time but remained infrequent compared to mTECI. mTECIII/IV largely appeared after birth (Fig. 4A, left).

**Figure 4.**
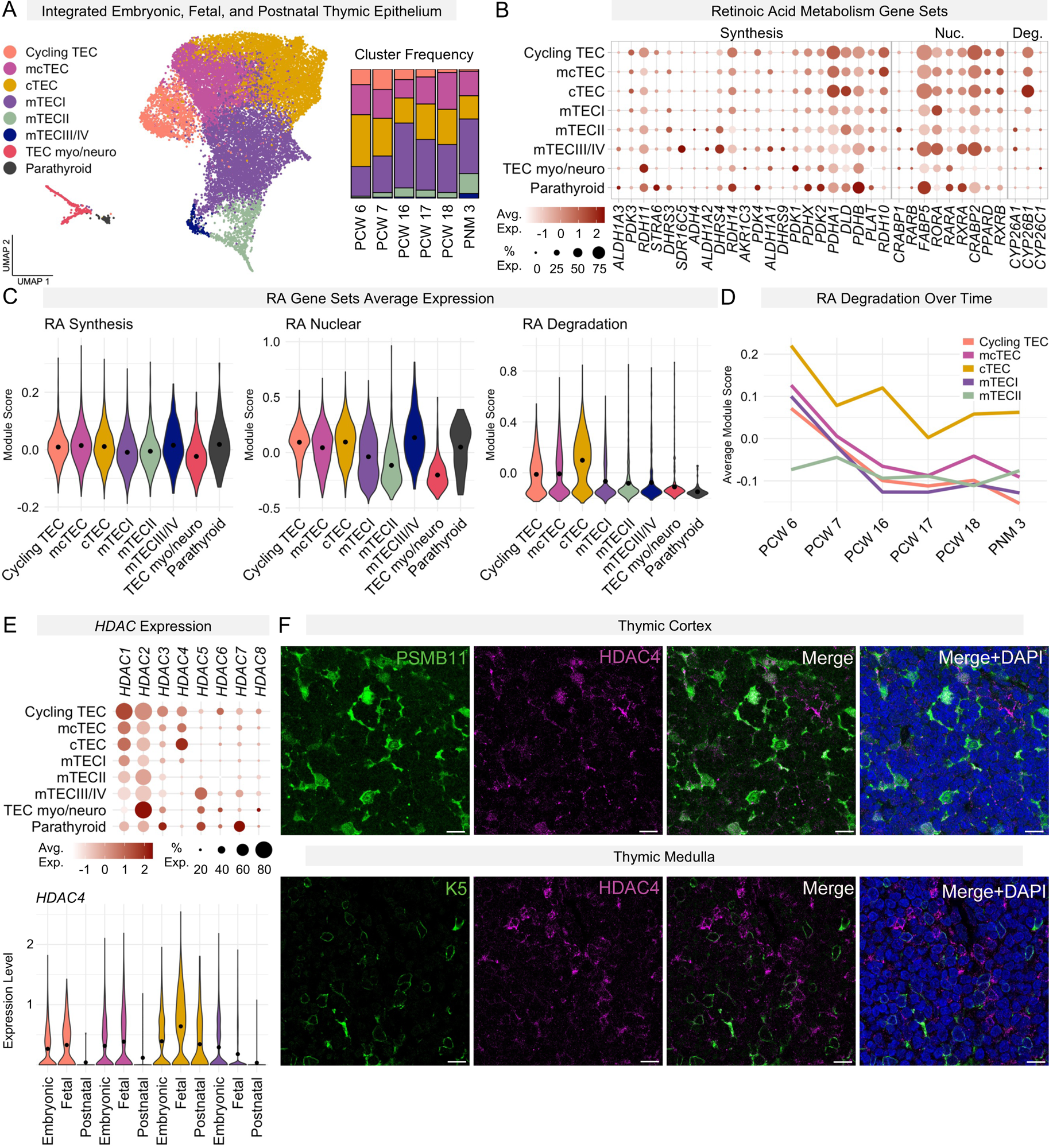
Cellular composition of the human thymus during development. **(A)** UMAP visualization of integrated TECs across all sampled time points, colored by developmental/postnatal age (left). Relative frequency of cells from select TEC subsets across advancing developmental ages (right). **(B)** Dot plot of gene sets defining retinoic acid synthesis, nuclear signaling, and degradation across select TEC subsets (Nuc, nuclear; Deg, degradation). **(C)** Violin plot representing the average expression of retinoic acid metabolism gene sets from (B) across visualized cell types from (A). RA, retinoic acid. **(D)** Average expression of retinoic acid metabolism gene sets across embryonic, fetal, and postnatal time points for select TEC subsets. **(E)** Dot plot of *HDAC 1-8* gene expression across thymic and parathyroid epithelial cell subpopulations (top). Violin plot of *HDAC4* gene expression in select TEC subsets across advancing developmental ages. Embryonic: pcw 6-7, Fetal: pcw 16-18, Postnatal: 3 months post birth (bottom). **(F)** Representative immunohistochemistry images of human fetal thymus sections stained for PSMB11 (green) and HDAC4 (magenta) protein. Counterstain with DAPI (blue). Scale bars: 50um.

Retinoic acid signaling plays an important role in vertebrate morphogenesis, specifically in the patterning of the pharyngeal pouches^61–63^. Moreover, the retinoic acid and IFN signaling axes intersect, with retinoic acid augmenting the effect of IFN^64^. To investigate differences in retinoic acid metabolism in thymic epithelial subsets, we integrated a curated set of developmentally relevant retinoic acid genes^65,66^ into expression modules for (I) retinoic acid synthesis, (II) retinoic acid nuclear receptor activity and (III) retinoic acid degradation (Fig. 4B). There were no gross differences in the retinoic acid synthetic capacity across analyzed TEC subsets. Retinoic acid nuclear activity showed lowest expression in AIRE-positive mTECII but peaked in post-AIRE mTECIII/IV. cTEC had a considerably higher expression of the retinoic acid degradation module (Fig. 4C). Retinoic acid degradation in cTEC, mcTEC and mTECI declined substantially during transition from early to midgestation (Fig. 4D). These data suggest distinct TEC subsets differentially metabolize retinoic acid.

Histone acetyltransferases and histone deacetylases (HDACs) are regulators of chromatin remodeling that can alter IFNα- and IFNγ-inducible transcription^67,68^. Recently, Hdac3 has been described as master regulator of murine mTEC development^69^, yet little is known about HDAC specificity in human TECs. Therefore, we examined *HDAC1-8* in our dataset. *HDAC4* showed uniquely high expression in cTEC that peaked midgestation and persisted after birth (Fig. 4E). CyTEC showed highest expression of *HDAC1*; mcTEC showed a similar HDAC profile as cyTEC, albeit confined to a smaller fraction of cells. The most highly expressed HDACs in mTECIII/IV and TECmyo/neuro were *HDAC5* and *HDAC2,* respectively. To corroborate preferential expression of *HDAC4* in cTEC, we performed IHC on human fetal thymus. HDAC4 preferentially colocalized with the cTEC marker PSMB11 (Fig. 4F), supporting our transcriptional findings. Distinct retinoic acid metabolism and HDAC expression profiles in individual TEC subsets raise the possibility that diverse TEC populations process IFN signals differentially.

### Patterns of peripheral tissue antigen expression

Central tolerance is induced in the thymus through negative selection of autoreactive T cells^70,71^. Negative selection of T cells recognizing peripheral tissue antigens (PTAs) is enabled by the TF *AIRE*^72^. To investigate signatures of PTA expression and their relationship to *AIRE*, we subclustered the medullary subpopulations mTECII, mTECIII/IV, TECmyo and TECneuro within the *vertical* dataset. mTECII and mTECIII/IV separated distinctly from the TECmyo and TECneuro populations (Fig. 5A). Further increase in granularity delineated six populations, including immature *AIRE*^low^ mTECII (*CCL19*), cycling mTECII (*TOP2A*), mature *AIRE*^high^ mTECII, post-*AIRE* mTECIII/IV (*SPINK5*), muscle-like TEC (*ACTA1*) and neuroendocrine-like TEC (*BEX1*) (Fig. 5B). Mature *AIRE*^high^ mTECII and post-*AIRE* mTECIII/IV populations significantly increased in frequency throughout the observed developmental interval, while muscle and neuroendocrine-like TEC did not expand notably beyond midgestation (Fig. 5C and D). Maturation and specification of medullary TECs from mTECI to mature *AIRE*^high^ mTECII stages was marked by decreasing *CCL19* and *TP63* expression. *AIRE*^high^ mTECII cells exclusively co-expressed *AIRE* and *FEZF2*. mTECIII/IV were high in *SPINK5* and *IVL* indicating their post-*AIRE* provenance^19,42,73^. Defining signatures in post-*AIRE* TECs included *GRHL1*, *GRHL3* and *KRTDAP*, denoting the keratinocyte-like state of mTECIII, while *POU2F3* indicated the emergence of Tuft-like cells (mTECIV)^19,20,42^. Neuroendocrine-like TEC (*NEUROD1)*, and muscle-like TEC (*MYOG1)* exhibited minimal keratinization and absent *FOXN1* expression but were high in *EPCAM*, supporting their epithelial nature (Fig. 5E). To assess if the transcriptional signatures defining TECmyo and TECneuro represent stable cellular phenotypes or isolated, possibly promiscuous gene expression, we probed for transcription factor-target gene interactions using SCENIC. Muscle and neuroendocrine-like TECs showed full expression of muscle- (*MYOD1, MYOG1, MEF2A, MEF2C, TEAD4, MYF6*) and neuroendocrine-specific (*NEUROD1, NEUROD2, SOX4, SOX11, POU4F1, NPDC1, ZEB1*) regulons, respectively (Fig. 5F), while *AIRE*-positive and post-*AIRE* medullary TECs did not express comprehensive sets of peripheral tissue defining regulons. Mature *AIRE*^high^ mTECII displayed highest MHC class I and II expression (Fig. 5G), in line with their highest type II IFN module score (Fig. 3D), whereas muscle and neuroendocrine-like TECs expressed low levels of MHC class I and no MHC class II (Fig. 5G). These findings point to divergent patterns of PTA expression in discrete thymic epithelial cell subsets. Absent expression of MHC class II molecules combined with low expression IFN response regulons (Fig. 2D) in TECmyo and TECneuro point to an IFN-independent developmental trajectory.

**Figure 5.**
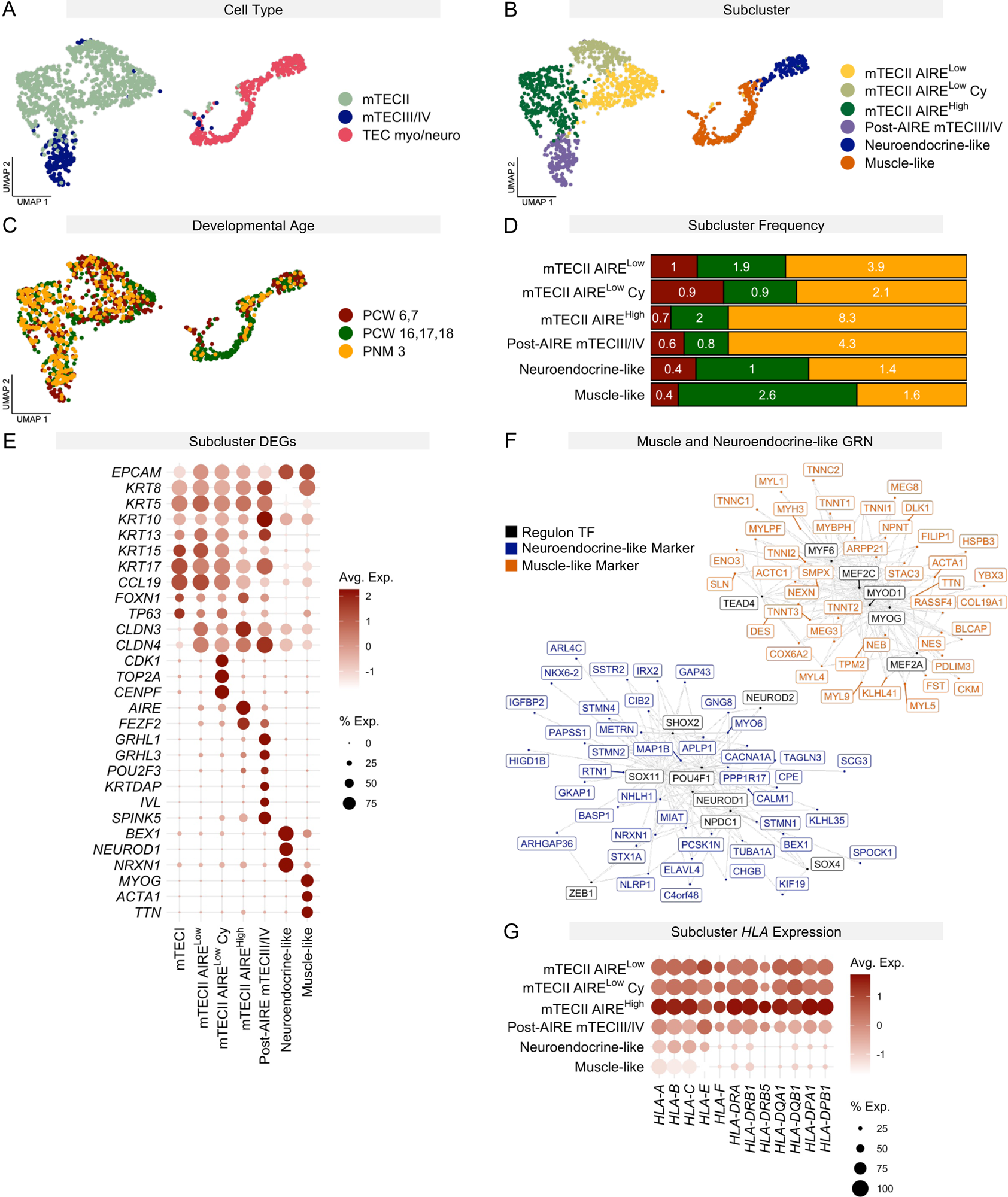
Expression patterns of rare TEC subsets. **(A)** UMAP visualization of mTECII, mTECIII/IV, and TECmyo/neuro across all developmental time points. Cells colored by cell type, **(B)** cells colored by higher clustering granularity, **(C)** cells colored by developmental age. **(D)** Stacked bar chart representing cluster frequency of mTECII, mTECIII/IV, and TECmyo/neuro at embryonic (red), fetal (green) and postnatal (yellow) stages related to Fig. 4A. **(E)** Dot plot of marker gene expression defining select subclusters. **(F)** Network visualization of highly expressed regulons with inferred interactions between regulon defining transcription factors and target genes in neuroendocrine- and muscle-like TECs. **(G)** Dot plot of select HLA gene expression across individual TEC subsets.

### Distinct IFN response networks direct cortical and medullary lineage specification

To dissect the transcriptional sequences that give rise to cortical and medullary thymic epithelial lineages across the *vertical* dataset, we employed tSpace, an algorithm optimized to elucidate developmental pathways from high-dimensional datasets^74^. The tSpace projection rendered an mcTEC-centric view of the thymus with cTEC and mTECI populations flanking mcTEC on both sides (Fig. 6A and fig. S10). Using Slingshot^75^ on the core populations cyTEC, mcTEC, cTEC and mTECI, we inferred two distinct trajectories along pseudotime, denoting cortical (trajectory 1) and medullary lineages (trajectory 2) (Fig. 6B). Notably, the trajectories originated at the cyTEC population but did not branch until midway through the mcTEC cluster.

**Figure 6.**
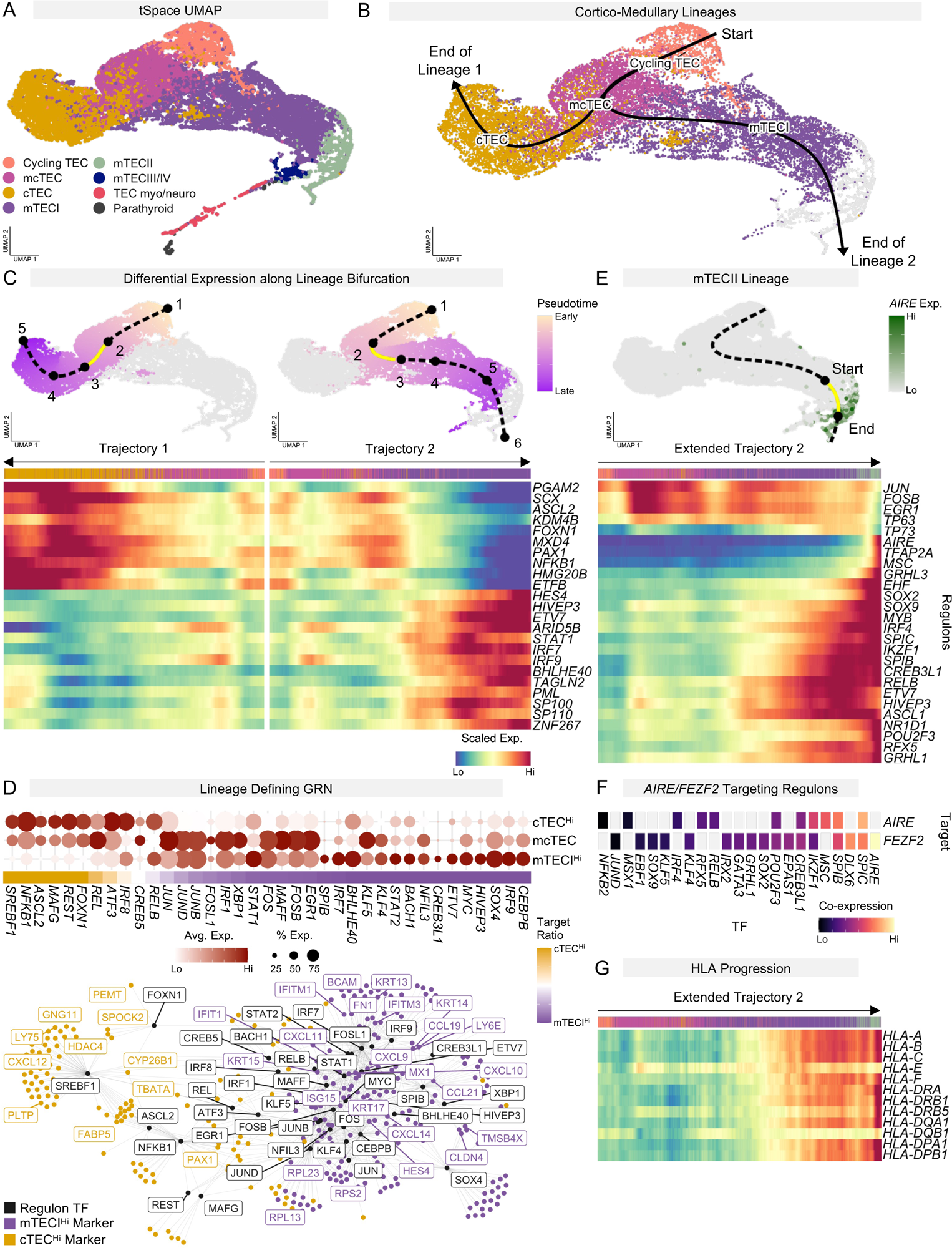
Regulatory networks defining cortical and medullary thymic epithelial lineages. **(A)** tSpace UMAP of TECs across advancing developmental and postnatal ages. Extended tSpace UMAP shown in supplementary figure S10. **(B)** Subset of tSpace UMAP from (A). Cortical and medullary lineage trajectories identified by Slingshot, plotted onto tSpace UMAP. **(C)** UMAP from (B) with cortical and medullary trajectories separately colored by pseudotime. Pseudotime intervals (knots) identified by TradeSeq are superimposed and numbered chronologically (top). Heatmaps representing the rolling average of select differentially expressed transcription factors between knots 2 and 3 along trajectories 1 (cortical) and 2 (medullary). From here onwards, heatmap color key corresponds to clusters in (A) (bottom). **(D)** Dot plot of regulon-defining transcription factors targeting lineage markers in populations with highest canonical lineage marker expression (cTEC^Hi^ and mTECI^Hi^) accompanied by heatmap representing the ratio of targets in the cTEC^Hi^ and mTECI^Hi^ populations, respectively (top). Network visualization of inferred interactions between regulon-defining transcription factors (black) and lineage markers (bottom). **(E)** tSpace UMAP from (B) with superimposed *AIRE* expression and extended medullary lineage trajectory. Highlighted pseudotime range corresponds to onset of *AIRE* expression (top). Heatmap representing the rolling average expression of select transcription factor regulons with differential expression within highlighted pseudotime range (bottom). **(F)** Heatmap of regulon-defining transcription factors targeting *AIRE* and *FEZF2.* **(G)** Heatmap of the rolling average expression of select *HLA* genes along the extended medullary lineage trajectory.

To explore transcriptional drivers directing the cortico-medullary lineage bifurcation, we investigated dynamic changes in gene expression along lineage trajectories using tradeSeq^76^. Gene expression was modeled along defined points (knots) in pseudotime. To uncover early signals that demarcate the cortico-medullary lineage branchpoint, we identified genes with differential expression across the immediate bifurcation (pseudotime region between knots 2 and 3). The key thymic regulators *PAX1* and *FOXN1*, together with *NFKB1* and *ASCL2,* demonstrated increased expression early along the cortical trajectory, whereas *IRF7*, *IRF9*, *STAT1,* the IFN-stimulated antigens *SP100*, *SP110*, and the immunomodulator *BHLHE40*^50^ showed an early medullary increase in expression (Fig. 6C and fig. S11).

To obtain a global view of the gene regulatory networks governing the divergent maturation of cTEC and mTECI, we applied SCENIC to populations with the highest canonical cortical (cTEC^Hi^) and medullary (mTECI^Hi^) lineage marker expression (fig. S12). Significant interactions between cTEC^Hi^ and mTEC^Hi^ markers and regulon-defining transcription factors were visualized in a network which formed distinct cortical and medullary clusters (Fig. 6D). In the cortical cluster, notable transcription factors with abundant target gene cross talk with cortex-defining markers included the basic helix-loop-helix-leucine zipper (bHLH-Zip) *SREBF1*, along with *FOXN1*, *NFKB1* and *ASCL2*, which shared interactions with *HDAC4*, the retinoic acid-binding and metabolizing proteins *FABP5* and *CYP26B1*, the thymic regulators *PAX1* and *TBATA* and other cTEC-defining markers such as *CXCL12* and *LY75*. Within the medullary cluster, *IRF1*, *IRF7, IRF9, STAT1* and *STAT2* shared abundant targets including the type II IFN response genes *CXCL9* and *CXCL10,* along with *CXCL11, CCL19, CCL21,* and multiple *KRT* genes. Upregulation of ribosomal proteins was unique to mTECI. To gain insights into the maturation of AIRE-positive TECs, we specifically evaluated the differential expression of regulons along the region of pseudotime corresponding to the onset of *AIRE* expression (Fig. 6E and fig. S13). The *AIRE*-targeting regulons *SPIB, SPIC, CREB3L1, IKZF1*, *POU2F3* and *RELB* exhibited a steady gain in expression along the extended medullary trajectory, followed by a brisk increase in *AIRE, TFAP2A* and *MSC* expression that marked the mTECII stage. This was paralleled by a sharp decline in *JUNB, FOSB, EGR1, TP63* and *TP73*. Notably, *SPIB, SPIC, CREB3L1, IKZF1* and *POU2F3* shared *FEZF2* and *AIRE* as target genes, while *FEZF2* emerged as *AIRE’s* most highly co-expressed target gene (Fig. 6F and fig. S14). MHC class I and II expression demonstrated consistent transcriptional gain along the medullary trajectory (Fig. 6G).

Using different computational models at varied points along pseudotime, *FOXN1, ASCL2*, *PAX1* and *NFκB1* consistently emerged as gene regulatory networks preferentially governing cortical lineage development while an activation of the interferon regulatory factor networks was predominantly observed in the medullary lineage. The expression of well-described signature genes defining cortical and medullary TEC validated the analysis.

## DISCUSSION

Progress in genomic analysis methods has redefined our concept of thymic epithelial architecture. The goal of this study was to understand the dynamic signals that drive thymic fate and lineage diversification during development. Unlike prior studies which solely focused on the thymus, we sought to gain novel insights by studying thymic development in an unprecedented context. Comparing human fetal epithelial cells from different anterior foregut-derived organs, we provide evidence that type I and II IFNs exert inductive signals on the thymic epithelial stroma. We further show that the pervasive effects of IFNs extend differentially into the thymic epithelial compartment, reflected in characteristic *NFκB* and *IRF* signatures specific to cortical and medullary lineages. Previous studies pointed to an activation of the TNF signaling axis, mediated by CD40 and RANK, during medullary TEC development^77,78^. These signals were shown to stimulate the NFκB pathway and proved essential for various aspects of mTEC fate and function^77,79,80^. Our data highlights a central role for type I and II IFNs during thymic morphogenesis with IRFs, STATs and NFκB emerging as downstream mediators that fine-tune IFN responses along cortical and medullary lineage trajectories. How exactly the global effects of type I and II IFNs are differentially mediated in cTECs and mTECs, is subject to ongoing investigations. Multiple lines of inquiry exist: Retinoic acid is known to increase the effect of type II IFN^64^. Type II IFN, in turn, has been shown to selectively increase the expression of *IRF1* and *STAT1*^81–83^. Our data indicates that retinoic acid is preferentially degraded in cTECs, which could explain the relatively lower average expression of IFN target genes in the cortex relative to the medulla. A similar rationale applies to HDACs as regulators IFN-inducible transcription^67,68^. Hence, we posit that differential expression patterns in genes altering the IFN response, along with other cytokines and their receptors, orchestrate spatially distinct niches within the thymus that ultimately give rise to diverse TEC subsets. However, this framework does not preclude that subtle differences in patterning underlying the thymic cortex and medulla precede the recruitment of IFN-releasing hematopoietic elements into the thymus and contribute to the higher baseline expression of *FOXN1*, *PAX1* and *ASCL2* in the cortex.

Starting at midgestation, the medullary lineage represents the largest fraction of TECs, highlighting its importance in negative selection. Negative selection must “anticipate” that T cells will encounter PTAs. This was originally believed to occur by cellular mimicry of medullary TECs resembling peripheral tissues^84^. The concept was largely abandoned with the discovery of the transcription factor Auto Immune Regulator (AIRE) in a subset of medullary TECs^72^. AIRE’s function is unique in that it triggers PTA expression independent of target-sequence specificity by leveraging chromatin plasticity^85^. FEZF2 has been described as an additional regulator of PTA expression^86^, although its mechanistic underpinnings remain less clear^87^. Together, AIRE and FEZF2 are thought to account for approximately 60% of PTA expression^134^, raising the question where the remaining PTA expression originates from. A recent study found that small numbers of mTECs with high PTA expression displayed distinct motif enrichments for lineage-defining transcription factors in a manner that was only partially Aire-dependent^42^. These findings led to the hypothesis that mTECs co-opt lineage defining transcription factors to induce tolerance^88^. We did not identify such motif enrichment in our human data set; however, our study suggests that AIRE-dependent PTA expression is an evolving process that may not yet be fully completed at birth. Of all queried TEC subsets, mature *AIRE^high^* mTECII showed the strongest IFNγ response signature, while IFN signaling did not seem to play a role in the development of TECmyo and TECneuro. In stark contrast to mature AIRE-positive TECs, TECmyo and TECneuro mimicked native muscle and neuroendocrine tissue signatures but lacked *FOXN1* expression, known post-AIRE markers, antigen presenting molecules and significant keratinization. These findings point to divergent patterns of AIRE-dependent and possibly AIRE-independent PTA expression in discrete thymic epithelial cell subsets.

Recently, a role for IL1b in the development of lung epithelial cells has been described^89^. Our work expands the emerging concept that local immune cell niches are essential for the development of different epithelial organs.

Between the extended maturation of the thymic medulla, i.e. mTECII/III/IV expand further after birth, and the involution of the thymus in the second decade of life, the window of thymic peak performance is short-lived. In addition, genetic defects and a myriad of iatrogenic and biological insults to the thymus can compromise its structural integrity and function. This underscores the urgent need for therapies to regenerate thymic tissues. Our study dissecting the transcriptional framework underlying commitment and specialization of the human thymic epithelial stroma during development reveals novel regulators of thymic epithelial differentiation and may offer actionable insights to advance the differentiation of thymic epithelial cells *in vitro*.

## MATERIALS AND METHODS

### Tissue Acquisition

Human fetal tissues from thymus, bronchus, lung and esophagus were obtained under Stanford School of Medicine IRB-approved protocols from the Department of Obstetrics and Gynecology at Stanford Hospital. Samples were derived from deidentified fetal tissue donations with written informed consent after elective termination of pregnancy. The study does not qualify as human subjects research, as defined by the Stanford School of Medicine IRB. Single cell transcriptomic data from pcw 6-7 samples was obtained in collaboration with the laboratory of Prof. Sarah Teichmann, from the MRC/Wellcome Trust-funded Human Developmental Biology Resources (HDBR, http://www.hdbr.org). Developmental age at embryonic stages up to pcw 8 was estimated using the Carnegie staging method (Molecular Genetics of Early Development, ISBN 1859960316). Sample C41 was originally estimated to be pcw 8 by the Carnegie staging method but was corrected to pcw 6 by comparison to other samples in the dataset. In fetal samples, developmental age was estimated from measurements of foot length and heel to knee length, compared against a standard growth chart^90^, and matched with the gestational age.

### Tissue Digestion

Human fetal thymus and lung samples (pcw 16 -18) were mechanically dissected into <2 mm^3^ pieces using surgical scissors. Human fetal bronchus and esophagus samples (pcw 16 -18) were opened longitudinally and mechanically scraped with scalpel to harvest the inner epithelial layer of bronchus and esophagus, respectively. All tissue samples were enzymatically digested with DNAse (10 mg/ml) and Liberase (2.5 mg/ml, Sigma Aldrich) per 10 g tissue using a tissue dissociator (gentleMACS, C tube, program m_spleen_02 (Miltenyi), 1-3 runs depending on tissue). Tissue fragments were transferred to a 50 ml tube and incubate at 37°C for 20 min. Cells were pelleted at 110 g, at RT for 2 min and resuspended in 10 ml PBS. For persisting fragments, the procedure was repeated 1-2 times to yield single cells. Cells were resuspended in MACS buffer (PBS, 1 % BSA, 2 mM EDTA), filtered through a 100 um cell strainer and counted.

### Epithelial Cell Enrichment

For fetal thymic tissue (16-18 pcw) only, cell suspension was depleted of thymocytes and other immune cells using Human CD45 MicroBeads (Miltenyi) and LD depletion columns (Miltenyi), 10^8^ cells/column, according to the manufacturer’s instructions. Single cell suspensions of all tissues, (CD45-depleted thymus, bronchus, lung, and esophagus) were enriched for epithelial cells using Human CD326 (EPCAM) Microbeads and LS positive selection columns (Miltenyi) according to the manufacturer’s instructions. Quality control for thymocyte depletion and epithelial cell enrichment for thymic single cell suspensions was performed by flow cytometry on FACSAria II with CD45-APC, CD3-FITC, CD205-PE, EPCAM-APC, and Live Dead-Near InfraRed-APC-Cy7 prior to MACS purification. Due to epitope blockage from magnetic bead antibodies, additional markers, i.e. CD7-AF647 (thymocytes) and CD105-PerCP/Cy5.5 (thymic mesenchymal cells) were used to assess post enrichment purity.

### scRNAseq Library Preparation and Sequencing

Gel-Bead in Emulsions (GEMs) were generated from single cell suspensions using the 3′ V3 chemistry and Chromium microfluidics system (10X Genomics). After reverse transcription, barcoded cDNA was extracted from the GEMs by Post-GEM RT-cleanup and amplified for 12 cycles. Amplified cDNA was subjected to 0.6x SPRI beads cleanup (Beckman, B23318). 25% of the amplified cDNA was subjected to enzymatic fragmentation, end-repair, A tailing, adaptor ligation and 10X specific sample indexing as per manufacturer’s protocol. Libraries were quantified using Bioanalyzer (Agilent) analysis. Libraries were pooled and sequenced on a NovaSeq 6000 instrument (Illumina) using the recommended sequencing read lengths of 28 bp (Read 1), 8 bp (i7 Index Read), and 91 bp (Read 2).

### scRNA-seq data processing

Single cell data were aligned to human genome library. After genome alignment and filtering for mitochondrial content and number of genes/reads per cell, the data was normalized with regularized negative Binomial regression using Seurat V.4.0^91^. Cells from different samples were then combined using integration anchors^92^. The dataset was organized by performing PCA dimensionality reduction on the 3000 most variable genes followed by unsupervised Louvain clustering on the 50 top principal components and visualized using Seurat’s UMAP function.

### Cumulative (bulk) RNA-seq Preparation and Sequencing

Total RNA was extracted from thymocytes and EPCAM positive-selected epithelial cells from thymus, bronchus and lung samples using the Qiagen RNeasy kit. Libraries were prepared from Total RNA using the SMART-Seq v4 Ultra low Input RNA kit (Takara Bio) and quantified using Bioanalyzer (Agilent) analysis. Libraries were pooled and sequenced on a NovaSeq 6000 instrument (Illumina) at 20 million paired reads per sample.

### Cumulative (bulk) RNA-seq processing and analysis

Bulk RNA-seq FASTQ data were processed through latch.bio, a cloud data platform to provide computing infrastructure and visualization tools for biological data analysis. The resulting count matrix was analyzed using the standard DESeq2/1.36.0 workflow. Normalized gene counts were visualized using ggplot2/3.40.

### Differential gene expression

Differential gene expression between single-cell clusters was determined using the FindAllMarkers function within Seurat.

### Single Cell Heatmap

The top 60 differentially expressed genes (DEGs) specific to fetal thymic epithelial cells compared to fetal epithelial clusters of all other organs were identified (*p*<10^-300). Downsampled cluster expression (30 cells per cluster using R’s sample function) of DEGs was visualized using Seurat’s DoHeatmap function.

### Gene set analysis

Gene sets were obtained from the Molecular Signature Database (MSigDB) curated C2 dataset^58^. We then performed gene set enrichment analysis (GSEA) using these sets and differential gene expression between annotated single-cell clusters as determined in Seurat. GSEA was performed with clusterProfiler/4.5.1.902 using the GSEA function (*p*<0.05). In order to visualize biological themes, we created a network representation of positively enriched gene set expression across single-cell clusters. In these networks, gene sets represent nodes and edges are drawn between nodes based on the proportion of shared genes between them (Jaccard similarity coefficient = 0.2). Each node is then colored proportionally by the relative enrichment of that gene set in each single-cell cluster. Gene set network visualization was performed using enrichplot/1.17.0.992^59^.

### Module scores

We determined gene module scores by applying the AddModuleScore function within Seurat to gene lists of interest.

### Gene regulatory network inference

The single cell regulatory network inference and clustering algorithm, SCENIC, was used to identify transcription factor (TF) regulons as previously described^47, 48^. Prior to pySCENIC/0.12.0 input (Python/3.9.0), log1p normalized expression matrices were filtered to remove low quality genes according to an average expression threshold value across all cells of 0.01.

In brief, sets of genes that are co-expressed with TFs are identified during SCENIC’s first step using the GRNBoost2^93^ algorithm. Then, each potential TF-target interaction was subjected to *cis*-regulatory motif analysis to detect putative direct-binding targets during the RcisTarget step. Interactions with significant motif enrichment of the respective upstream TF were retained while indirect targets lacking motif support were removed. Lastly, predicted regulons and an aggregate of their targets from all potential motifs were used to determine cellular enrichment for each regulon during the AUCell step.

### SCENIC AUCell Seurat Integration

SCENIC’s AUCell output matrix was passed into Seurat/4.3 and normalized across features using Seurat’s centered log ratio transformation. The data was subjected to a standard Seurat processing pipeline consisting of FindVariableFeatures(), ScaleData(), followed by RunPCA(). UMAP dimensionality reduction was carried out using the top 15 principal components for fetal anterior foregut epithelial cells. Differential regulons were identified with a log2fc.threshold = 0.01.

### SCENIC Network Visualization

The network visualization found in Figure 2C was made using the RcisTarget output of fetal anterior foregut epithelial cells. An adjacency matrix was constructed by drawing an edge if a regulon-defining TF was the target of another regulon-defining TF. To filter the network for the most highly co-expressed interactions, only interactions with a GRNBoost2 importance score >10 were retained (970 interactions out of a total of 7522). The resultant edge list was then passed into igraph/1.3.5 and clustered using igraph’s cluster_leiden function (resolution = 1). The network visualization found in Figure 6D was made using the RcisTarget output of cTEC and mTECI cells from all developmental stages with high canonical marker expression (*PRSS16* and *CCL19* log1p normalized expression value > 4). Seurat’s FindMarkers function was used to identify DEGs between populations (400 marker genes per population, *p*<10^-30). An adjacency matrix was constructed by drawing an edge if a regulon defining TF targeted a marker gene.

Highly co-expressed interactions were retained using a GRNBoost2 importance score >5 (1027 interactions out of a total of 5763). Only regulons with a minimum of 5 interactions were visualized. The network visualization found in Figure 5F was made using the RcisTarget output of mTECII, mTECIII/IV, and TEC myo/neuro cells from all developmental stages. Seurat’s FindMarkers function was used to identify DEGs between muscle-like and neuroendocrine-like clusters (*p*<10^-10). If a regulon-defining TF had >= 15 targets that intersected with the top 50 markers of either population, an edge was drawn. Highly co-expressed interactions were retained using a GRNBoost2 importance score >5 (300 interactions out of a total of 417). Network visualization of all networks was done using igraph/1.3.5, ggplot2/3.40, and ggnetwork/0.5.12.

### Trajectory and pseudotime analysis

We utilized tSpace/0.1.0 ^60^ to represent continuous relationships between single-cell expression profiles. In brief, tSpace determines cellular distances based on the shortest path between cellular k-nearest neighbor (KNN) subgraphs using the Wanderlust algorithm. These distances were then reduced using uniform manifold approximation and projection (UMAP). We then fit principal curves through dimensionally reduced tSpace data using slingshot/1.8.0 and SingleCellExperiment/1.12.0. Cycling TEC were selected as the starting-point cluster, while end-point clusters were not specified. We then extracted principal curve coordinates and cellular pseudotime values based on the resulting principal curves.

### Modeling gene expression along pseudotime

We utilized tradeSeq/1.4.0^61^ to build a negative binomial generalized additive model of single-cell gene expression and SCENIC AUCell values along pseudotime. This allowed for statistical testing of both within-trajectory and between-trajectory gene expression differences as well as smoothing of gene expression along pseudotime for the purpose of visualization. The earlyDETest function was used to identify the key differential TFs between lineages 1 and 2 across knots 2-3. The 40 TFs with the highest Wald Statistic value (*p*<0.05) were visualized. Tradeseq’s startVsEndTest function was used to identify within-lineage differentially expressed regulons along a pseudotime range corresponding to the onset of *AIRE* expression (medullary lineage, pseudotime range 22-28). The pseudotime range corresponding to the onset of *AIRE* expression was determined visually (Figure S14B). Regulons with a *p*<0.05 were visualized.

### Trajectory Heatmap Construction

Scaled expression values (Seurat’s ScaleData function) and quantile scaled (SCORPIUS/1.0.8 scale_quantile function) SCENIC Regulon AUC values were used to construct expression matrices along Slingshot pseudotime. The rolling average of these matrices (zoo/1.8-11) were then passed into pheatmap/1.0.12 for visualization.

### Co-expression Heatmap Construction

Inferred interactions between a regulon-defining TF and its target were identified using SCENIC’s RcisTarget output. These interactions were visualized according to co-expression values (GRNBoost2’s importance score) using ggplot2/3.40.

### Immunohistochemistry and confocal imaging

Paraffin-embedded human fetal thymus tissues were deparaffinized in xylene and rehydrated with ethanol solutions of decreasing concentrations until a final wash step with milliQ water. Antigen retrieval was performed with IHC-Tek^TM^ Epitope Retrieval Solution (#IW-1100-1L, IHC World) using steam. Tissue slices were permeabilized with permeabilization solution (PBS, 0.2%TritonX-100) for 30 minutes at room temperature, blocked with PBS containing 10% of Donkey Serum (DS) and 1% Bovine Serum Albumin for 1 hour at room temperature, and incubated with primary antibodies (see table 1 for reference) diluted in PBS with 1% DS in a humid chamber for 16 hours at 4°C. Slides were washed with PBS and incubated with fluorescently-labeled secondary antibody for 1 hour at room temperature. After subsequent washes with PBS, tissue autofluorescence was quenched with Vector TrueVIEW Autofluorescence Quenching Kit (# SP-8400, Vector Labs). Nuclei were stained with 5 µg/ml of DAPI in PBS for 10 minutes at room temperature. Slides were mounted with VectaMount. Tissue visualization and image collection was performed using a Leica Stellaris 5 Confocal microscope and LAS X software (Leica Microsystems). All images were acquired using 40X (HC PL APO CS2, NA 1.30) or 63X (HC PL APO CS2, NA 1.40) oil-immersion objectives.

**Table 1.** List of Antibodies.

| Antibody | Dilution | Source | Identifier |
| --- | --- | --- | --- |
| Mouse anti-human Histone deacetylase 4 (HDAC4) | 1:50 | Santa Cruz | sc-46671 |
| Mouse anti-rabbit Pan-Keratin | 1:100 | Santa Cruz | sc-81714 |
| Rabbit anti-human PSMB11 | 1:50 | Abcam | ab170832 |
| Rabbit anti-human Keratin 5 | 1:100 | Biolegend | 905501 |
| Donkey anti-rabbit Alexa Fluor 488 | 1:500 | Abcam | ab150073 |
| Donkey anti-mouse Alexa Fluor 647 | 1:500 | Abcam | ab150107 |

## ACKNOWLEDGEMENTS

We thank all members of Stanford 22q11 Deletion Syndrome Program, its directors and advisory board members (Maria Grazia Roncarolo, Irving Weissman, Gay Crooks, Antonio Baldini, Matthew Proteus) and all consortium participants (Kenneth I. Weinberg, Elizabeth Illingworth, Rosa Bacchetta, Andrea Cipriani, Hui Gai) for insightful input and discussions. We thank Hui Gai and MedGenome, Inc. for scRNA library preparation and sequencing, respectively. We thank Riley Suhar and Priscila F. Slepicka for graphical illustrations.

## FUNDING

California Institute of Regenerative Medicine, DISC-2 11109

Emerson Collective 1263707

Baxter Foundation

Specifically, we thank the anonymous donors of the Stanford 22q11 Deletions Syndrome Program for their generous support of our work.

## AUTHOR CONTRIBUTIONS

A.M., A.G., and K.G.W. interpreted the data and wrote the manuscript. The experiments were designed, performed and analyzed by A.M., B.S., A.G., and K.G.W.. P.F.S, K.M.H., H.D.N., W.W., M.A., D.K.K., M.C. and C.Y.Y. performed additional experiments and bioinformatic data processing. V.W., S.T., J.P. and V.K. provided samples and analytical input. G.A.H., C. G., V.S., and P.K. reviewed design, bioinformatic analysis and edited the manuscript.

## DATA AVAILABILITY

Single cell RNA-seq data was deposited in GEO with accession number GSE212244. Bulk RNA-seq data was deposited in GEO with accession number GSE241479. All code is available at the following Github repository: https://github.com/abdulvasey/Thymic-Development-Figures. All other data needed to support the conclusions of the paper are present in the paper or the Supplementary Materials.

---

**Supplementary Figure S1.**
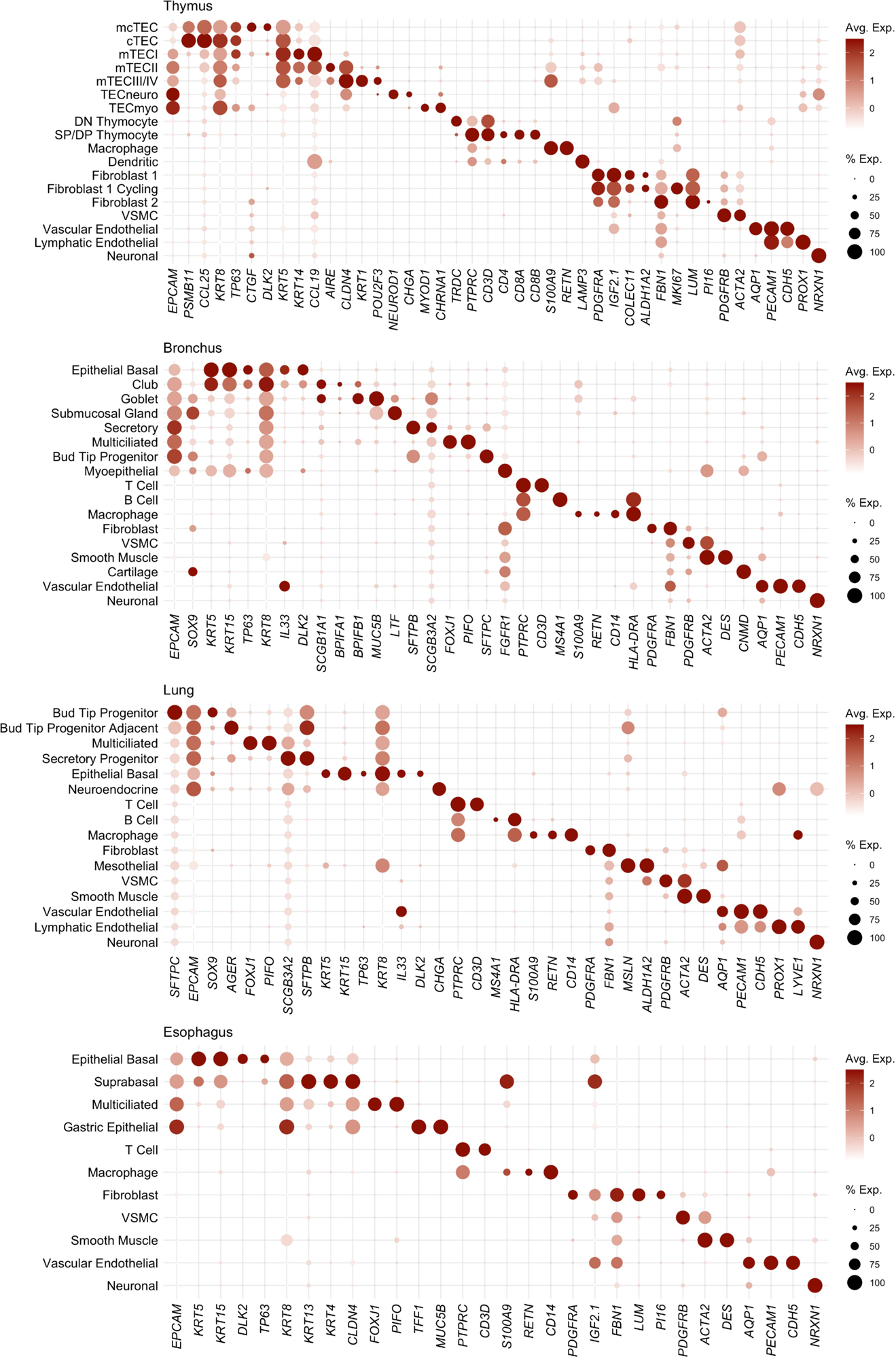
Dot plot of marker gene expression for annotated cell types in fetal thymus, bronchus, lung, and esophagus.

**Supplementary Figure S2.**
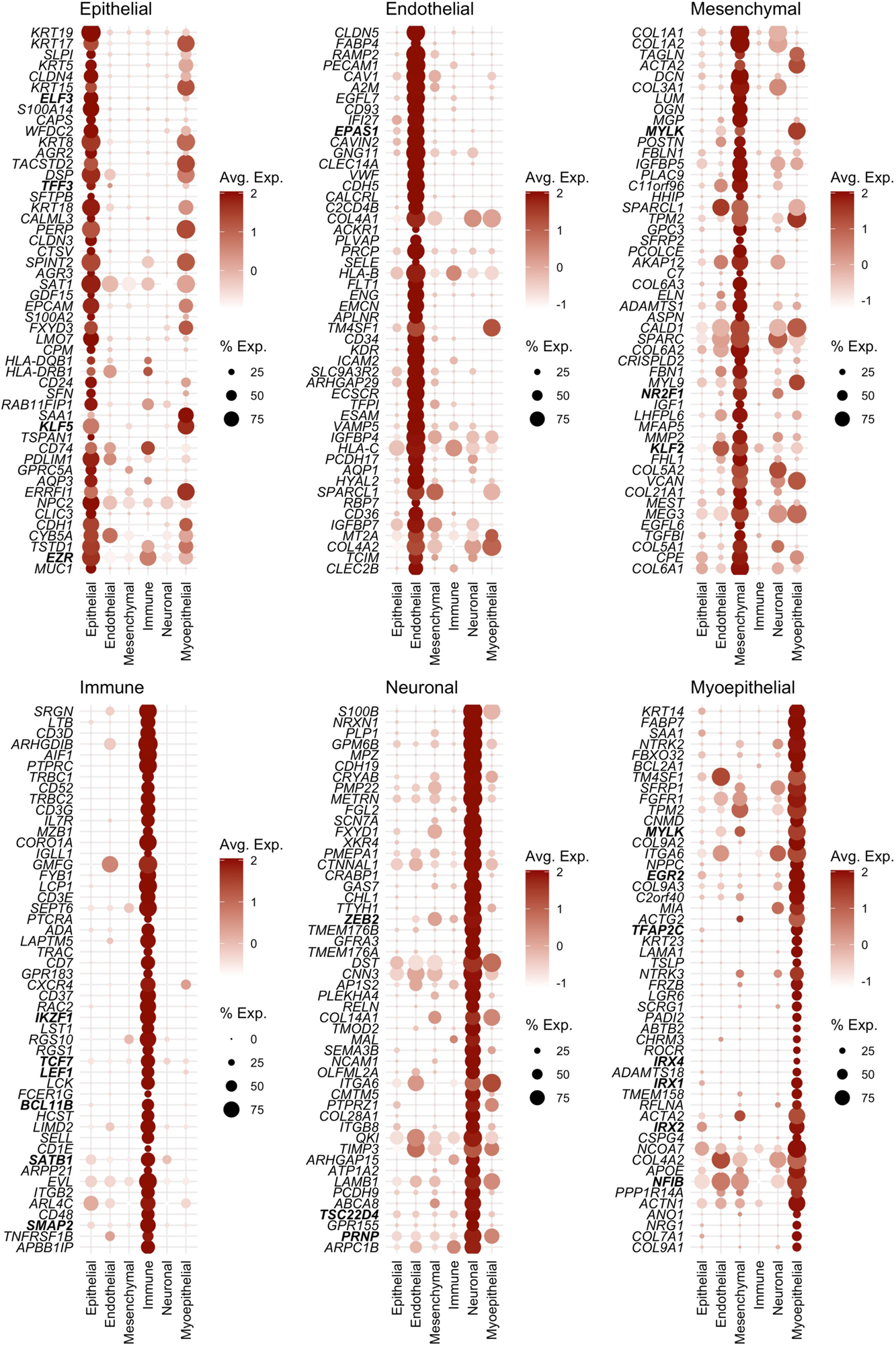
Dot plots of top differentially expressed genes for all major tissue types across cells from human fetal thymus, bronchus, lung and esophagus (transcription factors are labeled in bold).

**Supplementary Figure S3.**
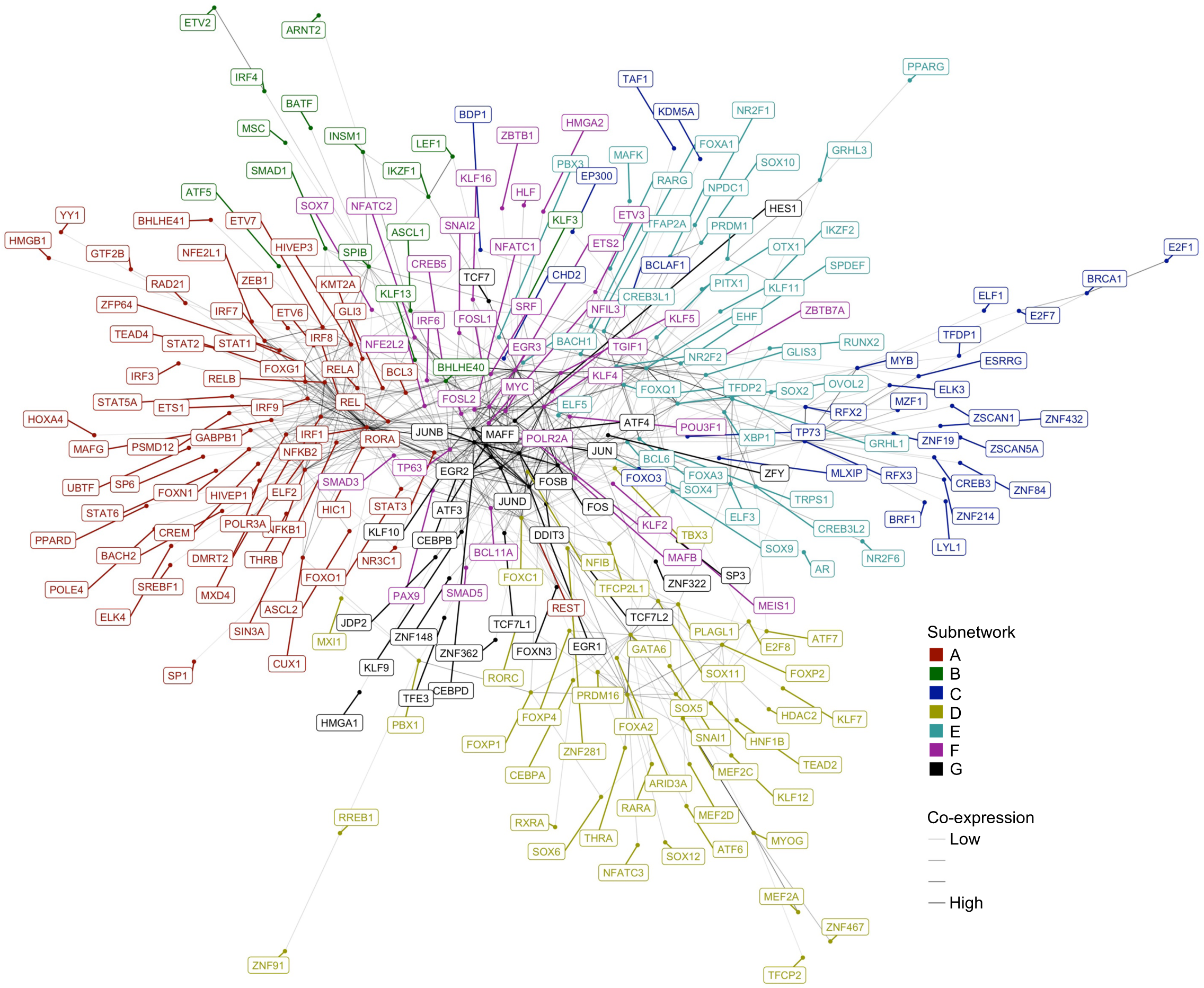
Fully annotated version of network from Fig. 2C. Visualization of transcription factor regulon subnetworks in all fetal epithelial cells defined by Leiden algorithm. Line thickness corresponds to co-expression of interaction.

**Supplementary Figure S4.**
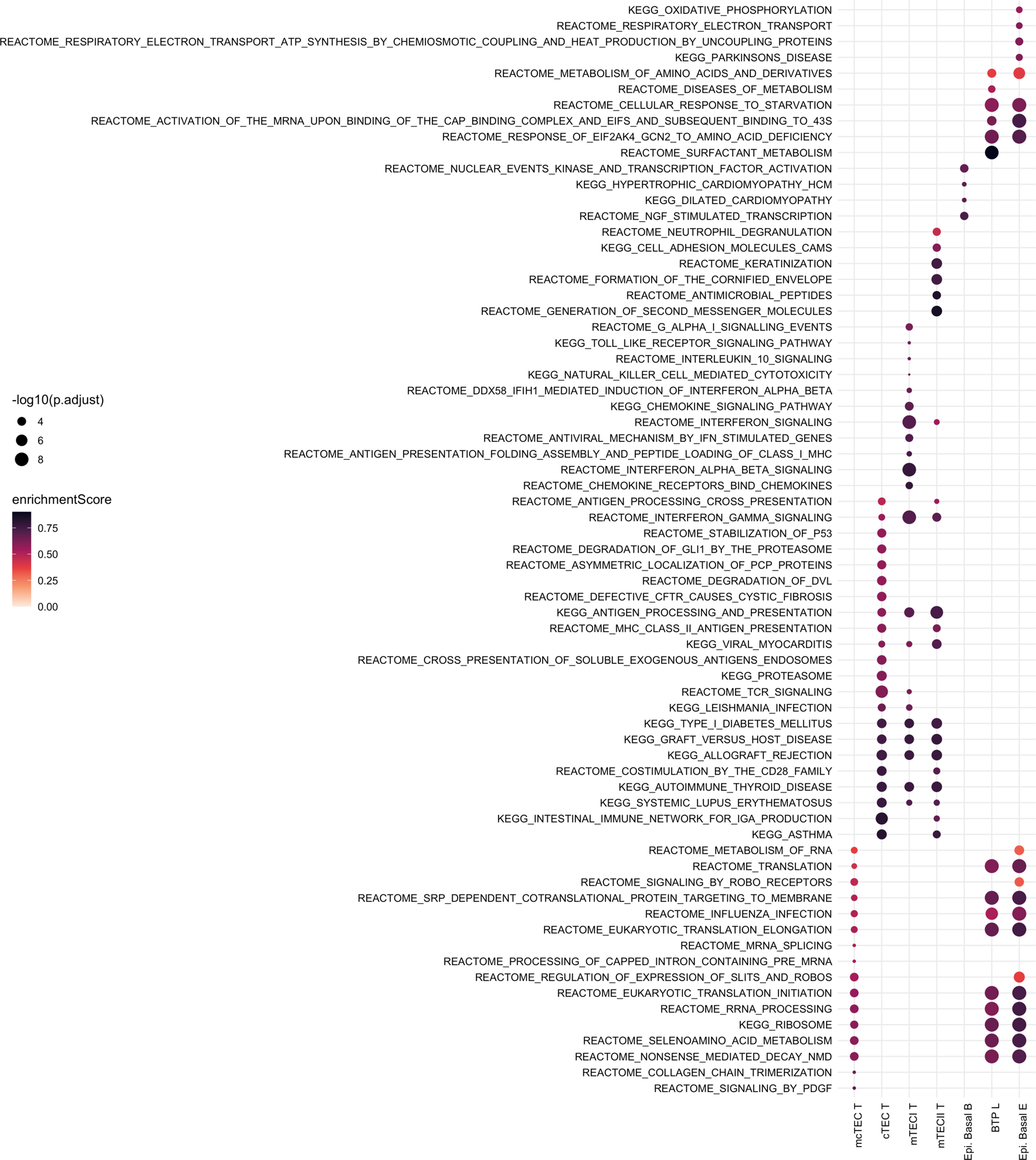
Dot plot of enriched gene sets across select epithelial cell subpopulations for thymus, bronchus, lung and esophagus at fetal stages of development. Dot size corresponds to p-value, color corresponds to enrichment score.

**Supplementary Figure S5.**
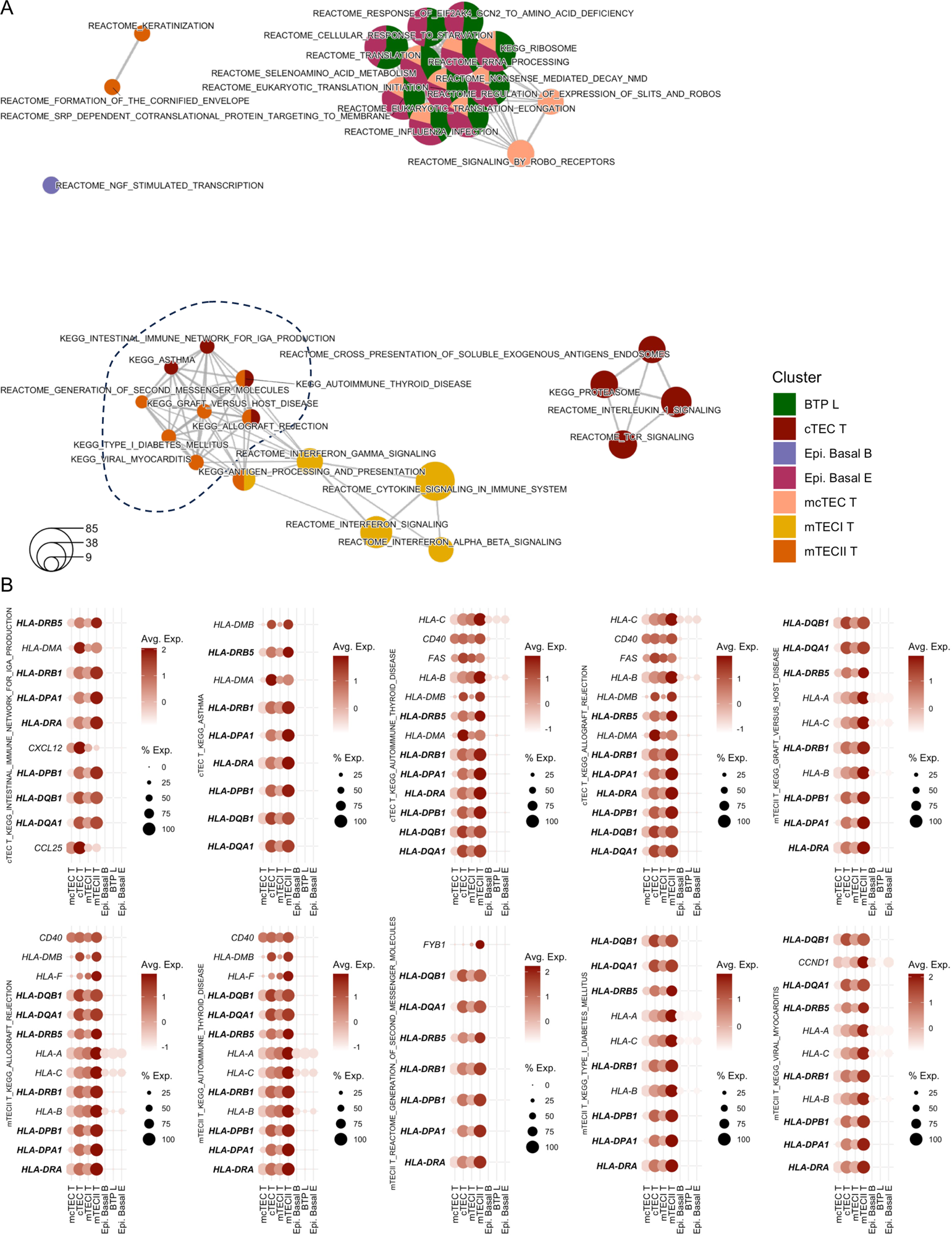
**(A)** Fully annotated version of GSEA network found from Fig. 3B. Network visualization of differentially expressed GSEA gene sets across select epithelial cell subpopulations for thymus, bronchus, lung and esophagus at fetal stages of development. Each node represents a specific GSEA gene set (*p*<0.05). Edges between nodes denote a significant number of shared genes between nodes. Node size reflects the number of genes within each gene set. **(B)** Enriched genes from the gene sets within the dotted circle are visualized in dot plots. Emboldened genes correspond to common genes across dot plots.

**Supplementary Figure S6.**
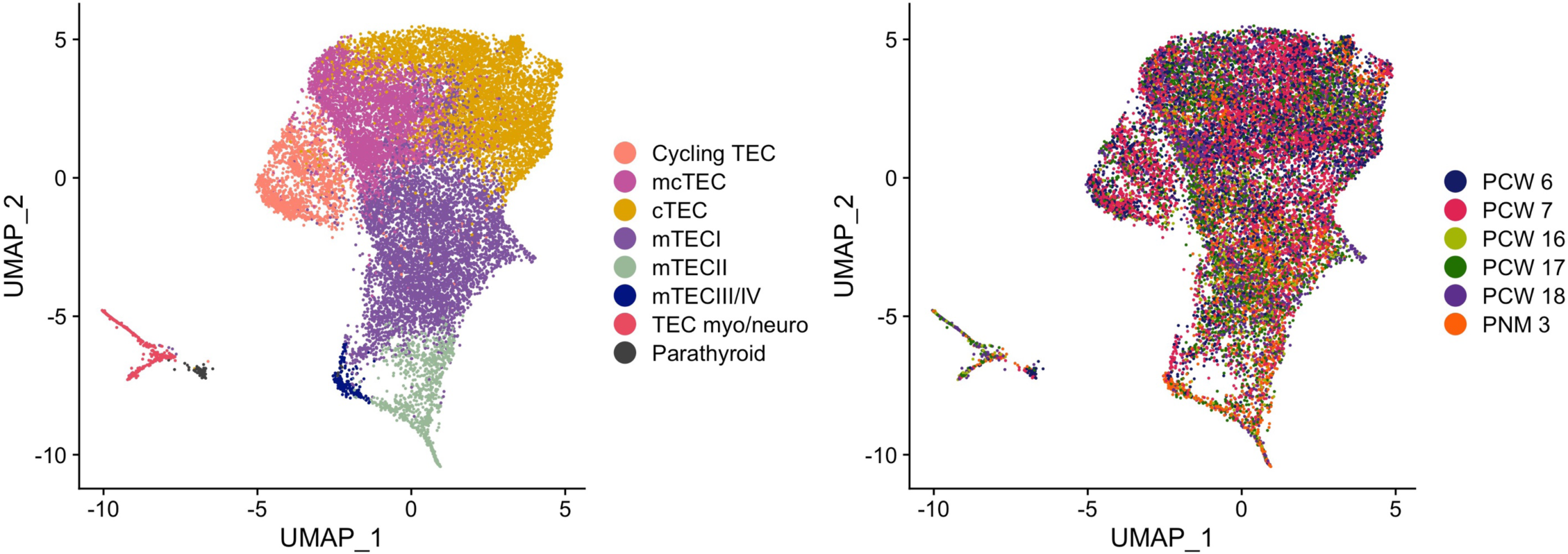
UMAP visualization of integrated TEC across all sampled time points, colored by developmental/postnatal age (left) and cell type (right).

**Supplementary Figure S7.**
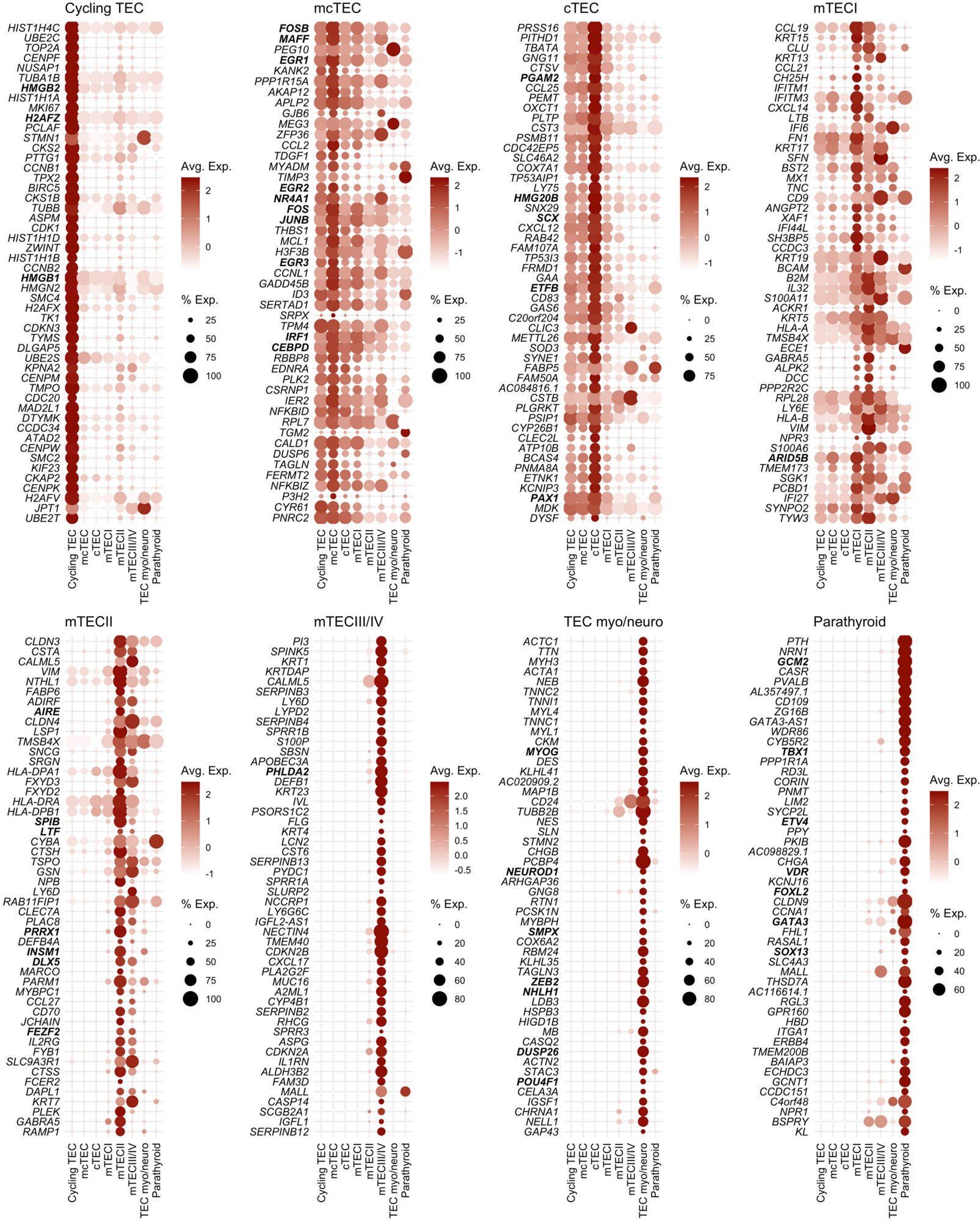
Dot plots of differentially expressed genes for thymic epithelial and parathyroid clusters across all sampled developmental time points (transcription factors are labeled in bold).

**Supplementary Figure S8.**
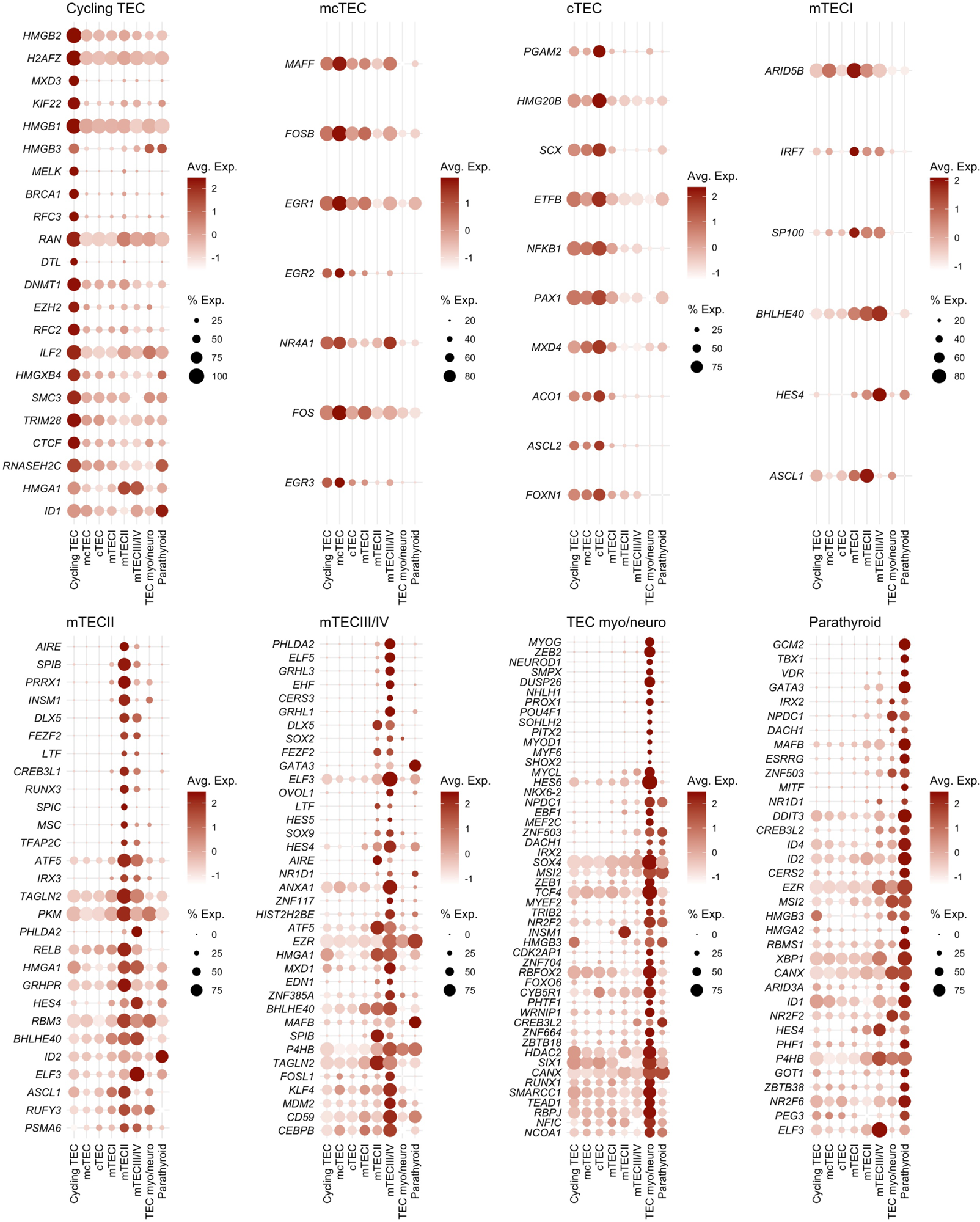
Dot plots of differentially expressed transcription factors for thymic epithelial and parathyroid clusters across all sampled developmental time points.

**Supplementary Figure S9.**
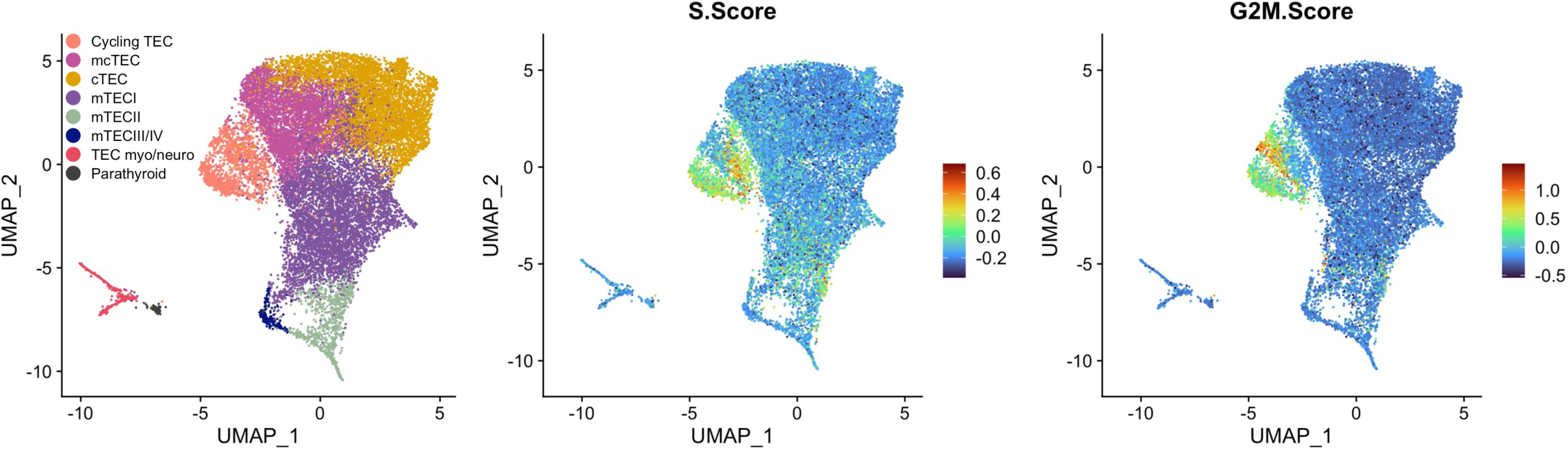
UMAP visualization of TECs across all sampled developmental time points colored by Seurat S and G2M phase scores.

**Supplementary Figure S10.**
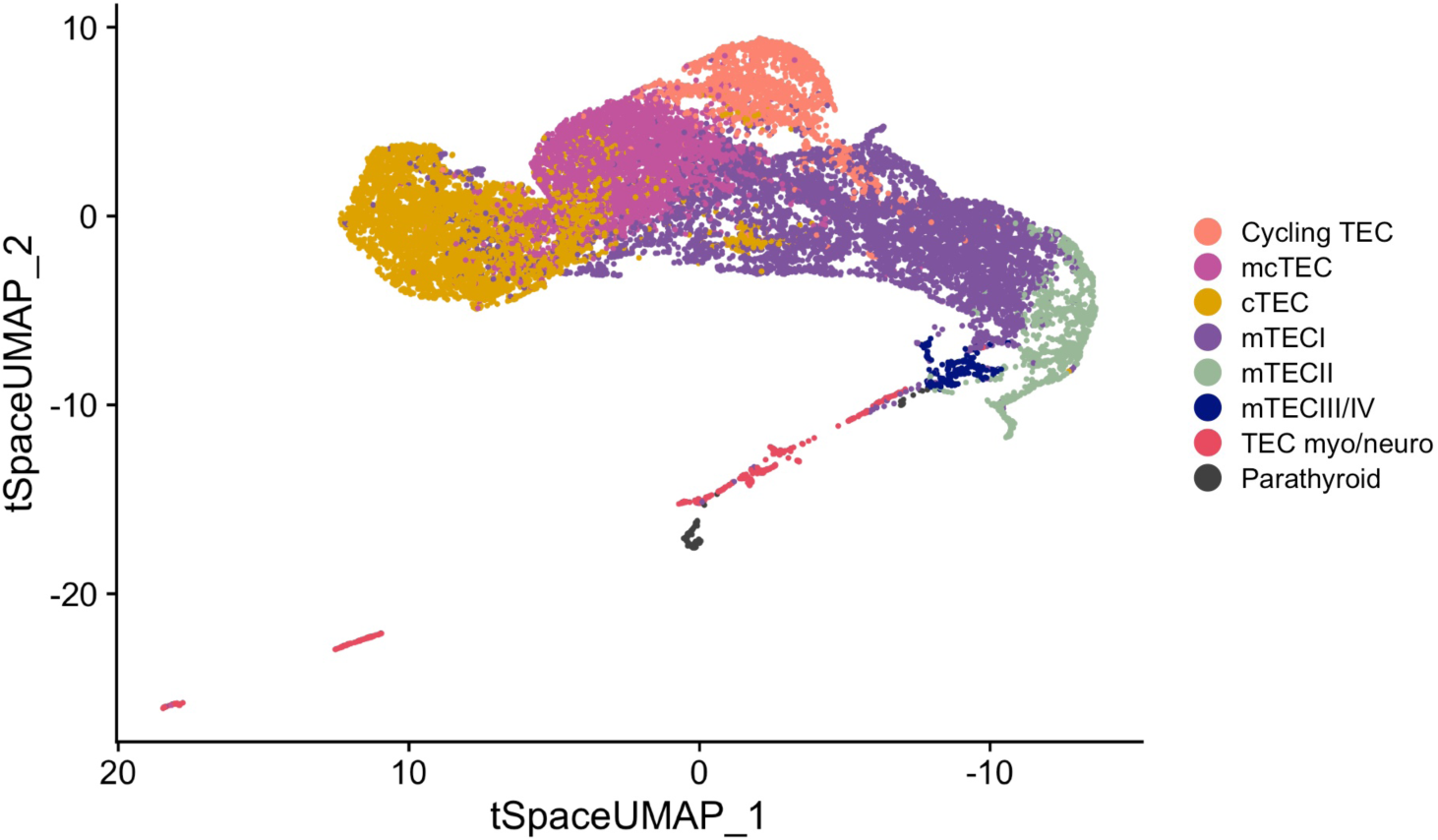
tSpace UMAP visualization of all integrated TECs from embryonic, fetal and postnatal stages of development.

**Supplementary Figure S11.**
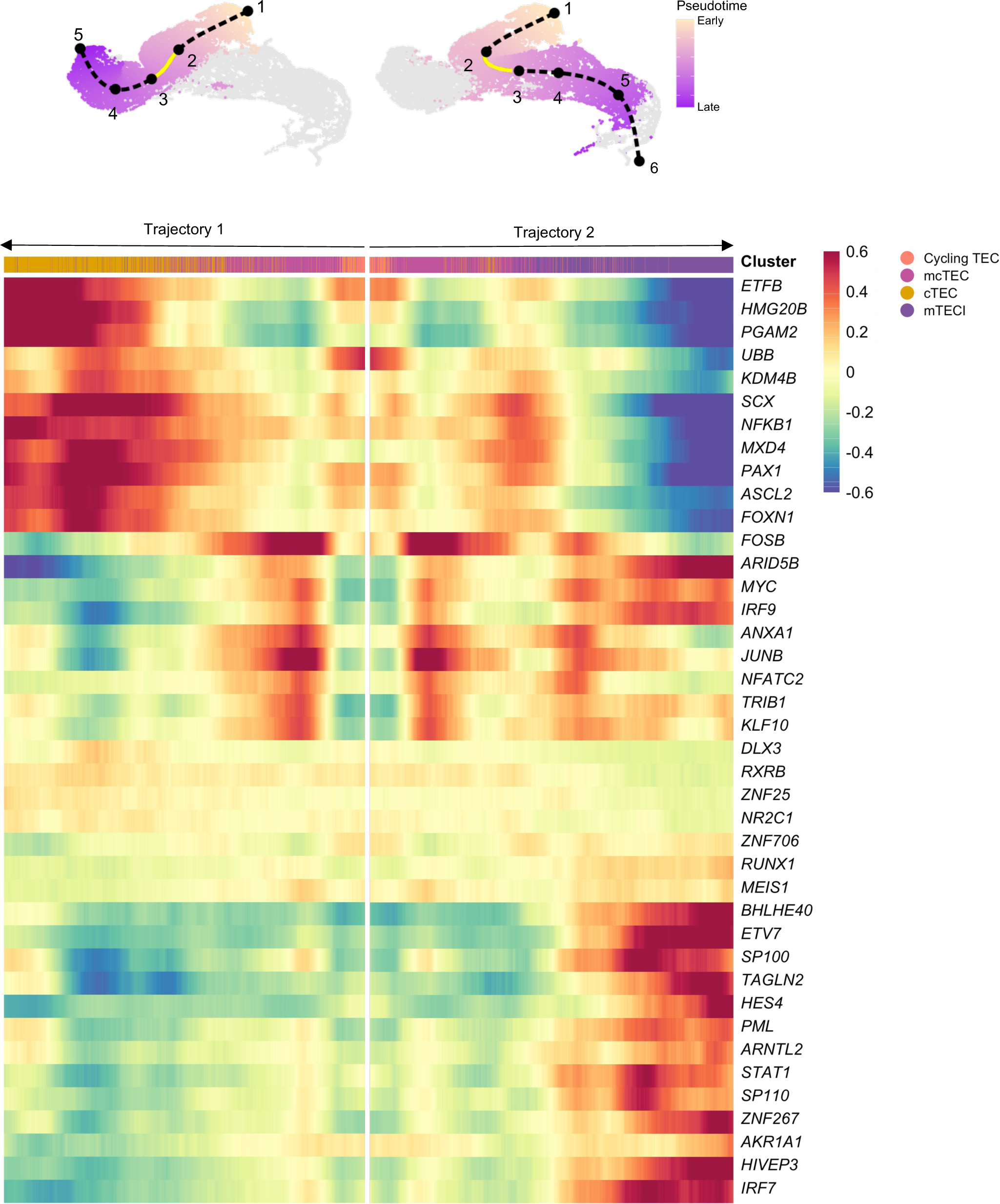
Subset of tSpace UMAP from Fig. 6B with cortical and medullary trajectories separately colored by pseudotime. Pseudotime intervals (knots) identified by TradeSeq are superimposed and numbered chronologically (top). Heatmaps representing the rolling average of the top 40 differentially expressed transcription factors between knots 2 and 3 along trajectories 1 and 2 (bottom).

**Supplementary Figure S12.**
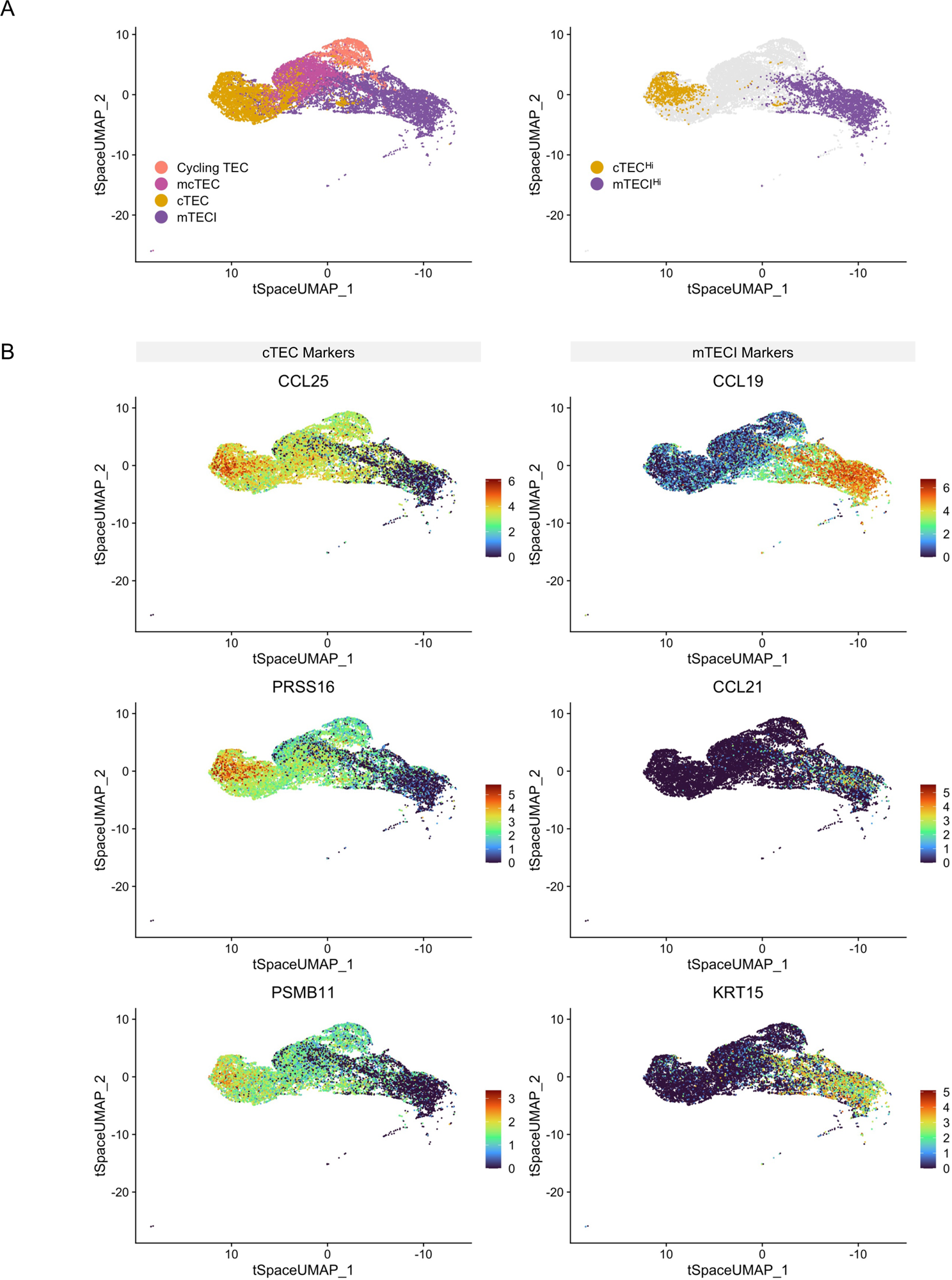
**(A)** tSpace UMAP visualization of integrated TECs from embryonic, fetal and postnatal stages of development. mTECII, mTECIII/IV, TECmyo/neuro, and parathyroid epithelial cells not visualized (left). tSpace UMAP highlighted by populations with high canonical lineage marker expression (right). **(B)** tSpace UMAP with superimposed canonical cTEC and mTECI marker expression.

**Supplementary Figure S13.**
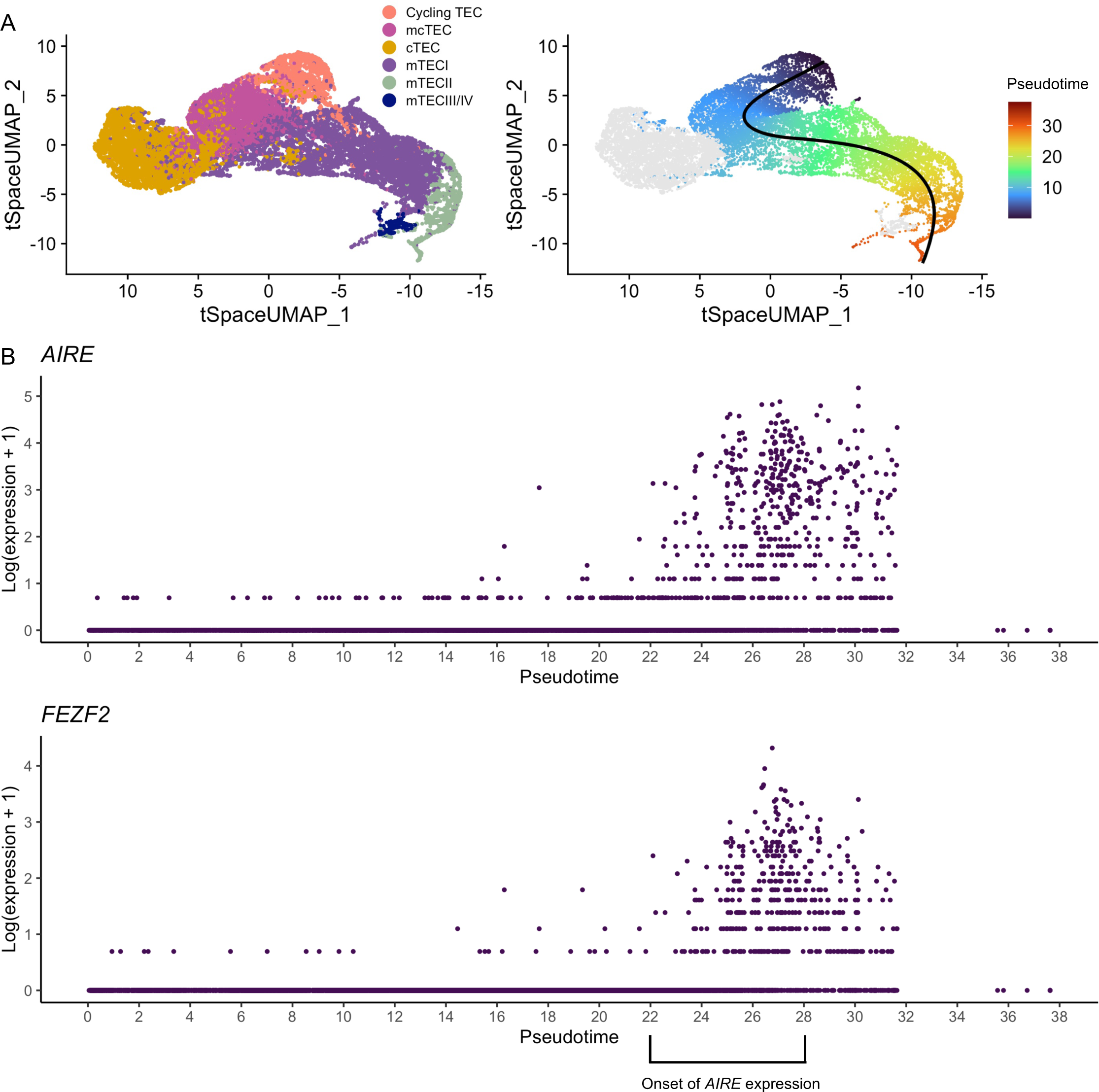
**(A)** tSpace UMAP from Fig. 6B colored by cell type (left) and pseudotime values along the extended medullary trajectory (right). **(B)** Gene expression of cells along pseudotime of the extended medullary trajectory.

**Supplementary Figure S14.**
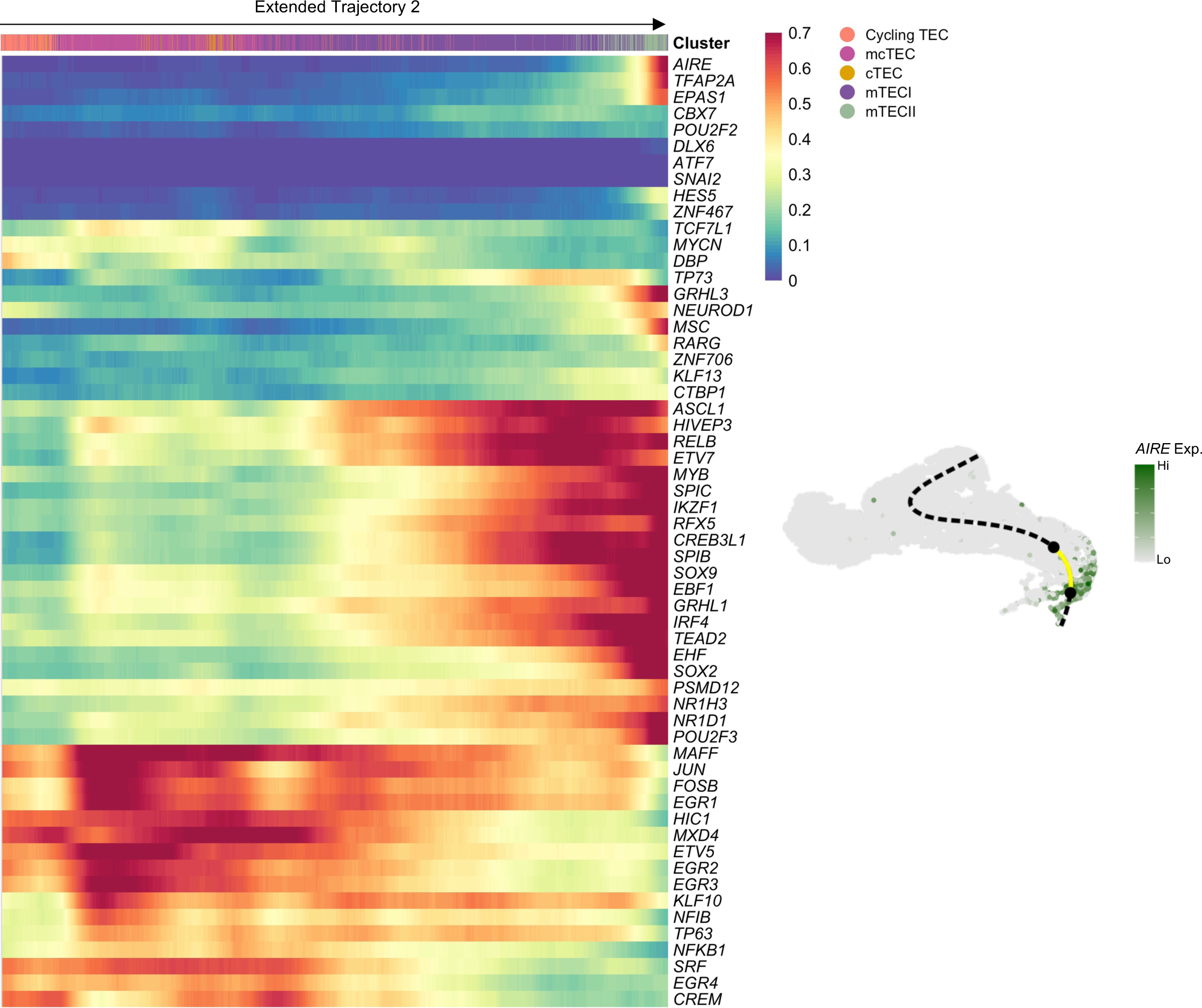
Heatmap representing the rolling average expression of all transcription factor regulons with differential expression (p<0.05) along highlighted pseudotime region superimposed on tSpace UMAP from Fig. 6B.

**Supplementary Table S1.** Sample metadata. [Sample ID] TH: Thymus, B: Bronchus, L: Lung, E: Esophagus

| Sample ID | Organ | Age | Number of Cells | Mean Counts Per Cell | Mean Genes Per Cell | Median Mito Percentage | Gender | Tissue processing |
| --- | --- | --- | --- | --- | --- | --- | --- | --- |
| C40 TH TOT 2 | Thymus | Post Conception Week 7 | 7180 | 11139 | 3054.5 | 1.97 | Female | All cells |
| C40 TH TOT 1 | Thymus | Post Conception Week 7 | 6841 | 11293.8 | 3098.1 | 1.97 | Female | All cells |
| C41 TH TOT 2 | Thymus | Post Conception Week 6 | 5789 | 14190.5 | 3544.9 | 2.61 | Male | All cells |
| C41 TH TOT 1 | Thymus | Post Conception Week 6 | 5620 | 14244.3 | 3547.6 | 2.52 | Male | All cells |
| KGW1 TH EPCAM 45 | Thymus | Post Conception Week 16 | 3046 | 21747 | 4135 | 7.29 | Male | EPCAM positive selection, CD45 depletion |
| F83 TH EPCAM | Thymus | Post Conception Week 17 | 3678 | 9575.2 | 2634.2 | 2.55 | Female | EPCAM FACS sorted |
| KGW2 TH EPCAM 45 | Thymus | Post Conception Week 17 | 7634 | 11754 | 2993 | 6.81 | Female | EPCAM positive selection, CD45 depletion |
| KGW3 TH EPCAM 45 | Thymus | Post Conception Week 18 | 8648 | 9349 | 2721 | 5.73 | Male | EPCAM positive selection, CD45 depletion |
| T07 TH EPCAM S3 | Thymus | 3 Months Postnatal | 3149 | 11355.6 | 2819.6 | 3.86 | Male | EPCAM FACS sorted |
| KGW4 B EPCAM | Bronchus | Post Conception Week 16 | 5920 | 15651 | 3525 | 7.69 | Male | EPCAM positive selection |
| KGW5 B EPCAM | Bronchus | Post Conception Week 17 | 4376 | 16449 | 3188 | 8.56 | Female | EPCAM positive selection |
| KGW6 B EPCAM | Bronchus | Post Conception Week 18 | 7414 | 15387 | 2979 | 9.43 | Male | EPCAM positive selection |
| KGW7 L EPCAM | Lung | Post Conception Week 16 | 3726 | 20366 | 3862 | 7.95 | Male | EPCAM positive selection |
| KGW8 L EPCAM | Lung | Post Conception Week 17 | 7075 | 11074 | 2810 | 8.11 | Female | EPCAM positive selection |
| KGW9 L EPCAM | Lung | Post Conception Week 18 | 7208 | 12965 | 3254 | 7.12 | Male | EPCAM positive selection |
| KGW10 E EPCAM | Esophagus | Post Conception Week 16 | 4603 | 16004 | 3513 | 7 | Male | EPCAM positive selection |
| KGW11 E EPCAM | Esophagus | Post Conception Week 17 | 6464 | 11431 | 2723 | 8.48 | Female | EPCAM positive selection |

**Supplementary Table S2.**
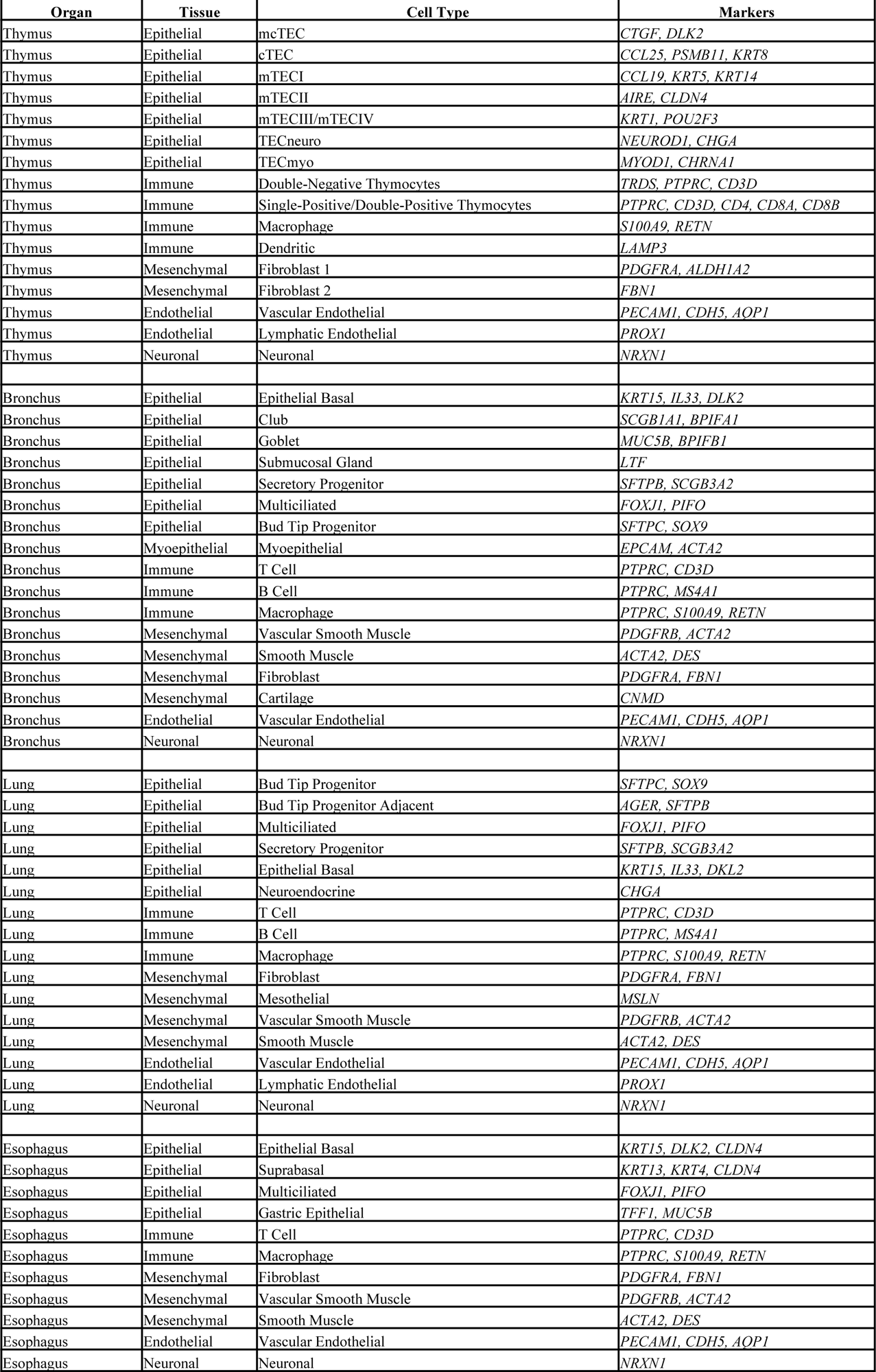
Table contains canonical markers used for cluster annotation.

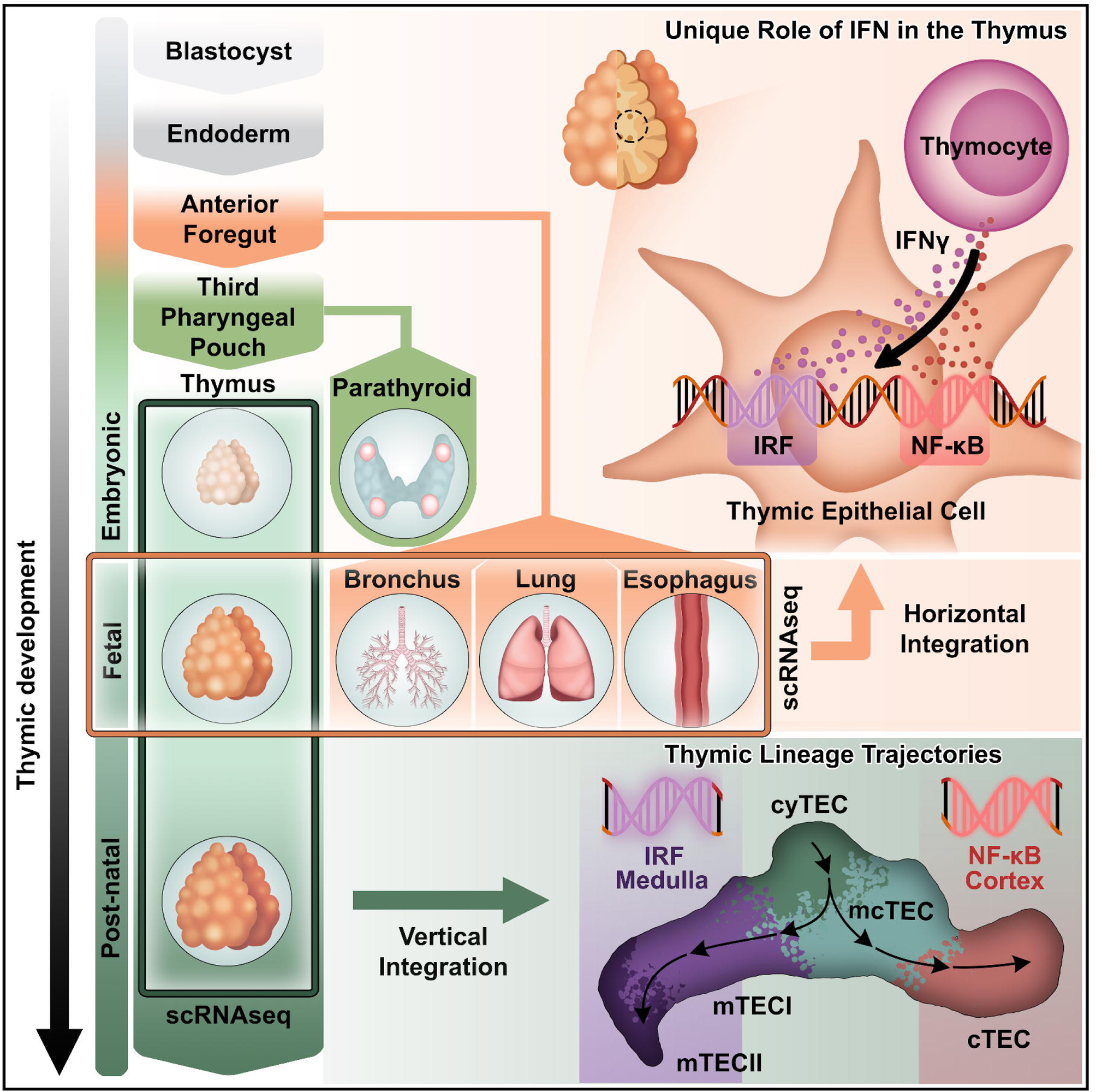

## REFERENCES

1. Miller JF, Osoba D. Current concepts of the immunological function of the thymus. Physiol Rev. Jul 1967;47(3):437–520. doi:10.1152/physrev.1967.47.3.437

2. Abramson J, Anderson G. Thymic Epithelial Cells. Annu Rev Immunol. Apr 26 2017;35:85–118. doi:10.1146/annurev-immunol-051116-052320

3. Kadouri N, Nevo S, Goldfarb Y, Abramson J. Thymic epithelial cell heterogeneity: TEC by TEC. Nat Rev Immunol. Apr 2020;20(4):239–253. doi:10.1038/s41577-019-0238-0

4. Miller JF. The golden anniversary of the thymus. Nat Rev Immunol. May 27 2011;11(7):489–95. doi:10.1038/nri2993

5. Palmer S, Albergante L, Blackburn CC, Newman TJ. Thymic involution and rising disease incidence with age. Proc Natl Acad Sci U S A. Feb 20 2018;115(8):1883–1888. doi:10.1073/pnas.1714478115

6. Hollander GA, Widmer B, Burakoff SJ. Loss of normal thymic repertoire selection and persistence of autoreactive T cells in graft vs host disease. J Immunol. Feb 15 1994;152(4):1609–17.

7. Haynes BF, Markert ML, Sempowski GD, Patel DD, Hale LP. The role of the thymus in immune reconstitution in aging, bone marrow transplantation, and HIV-1 infection. Annu Rev Immunol. 2000;18:529–60. doi:10.1146/annurev.immunol.18.1.529

8. Thomas R, Wang W, Su DM. Contributions of Age-Related Thymic Involution to Immunosenescence and Inflammaging. Immun Ageing. 2020;17:2. doi:10.1186/s12979-020-0173-8

9. McElhaney JE, Effros RB. Immunosenescence: what does it mean to health outcomes in older adults? Curr Opin Immunol. Aug 2009;21(4):418–24. doi:10.1016/j.coi.2009.05.023

10. Zorn AM, Wells JM. Vertebrate endoderm development and organ formation. Annu Rev Cell Dev Biol. 2009;25:221–51. doi:10.1146/annurev.cellbio.042308.113344

11. Graham A, Richardson J. Developmental and evolutionary origins of the pharyngeal apparatus. Evodevo. Oct 1 2012;3(1):24. doi:10.1186/2041-9139-3-24

12. Kishimoto K, Iwasawa K, Sorel A, et al. Directed differentiation of human pluripotent stem cells into diverse organ-specific mesenchyme of the digestive and respiratory systems. Nat Protoc. Aug 17 2022;doi:10.1038/s41596-022-00733-3

13. Green MD, Chen A, Nostro MC, et al. Generation of anterior foregut endoderm from human embryonic and induced pluripotent stem cells. Nat Biotechnol. Mar 2011;29(3):267–72. doi:10.1038/nbt.1788

14. Miller AJ, Dye BR, Ferrer-Torres D, et al. Generation of lung organoids from human pluripotent stem cells in vitro. Nat Protoc. Feb 2019;14(2):518–540. doi:10.1038/s41596-018-0104-8

15. Trisno SL, Philo KED, McCracken KW, et al. Esophageal Organoids from Human Pluripotent Stem Cells Delineate Sox2 Functions during Esophageal Specification. Cell Stem Cell. Oct 4 2018;23(4):501–515 e7. doi:10.1016/j.stem.2018.08.008

16. Zhang Y, Yang Y, Jiang M, et al. 3D Modeling of Esophageal Development using Human PSC-Derived Basal Progenitors Reveals a Critical Role for Notch Signaling. Cell Stem Cell. Oct 4 2018;23(4):516–529 e5. doi:10.1016/j.stem.2018.08.009

17. Huang SX, Green MD, de Carvalho AT, et al. The in vitro generation of lung and airway progenitor cells from human pluripotent stem cells. Nat Protoc. Mar 2015;10(3):413–25. doi:10.1038/nprot.2015.023

18. Kernfeld EM, Genga RMJ, Neherin K, Magaletta ME, Xu P, Maehr R. A Single-Cell Transcriptomic Atlas of Thymus Organogenesis Resolves Cell Types and Developmental Maturation. Immunity. Jun 19 2018;48(6):1258–1270 e6. doi:10.1016/j.immuni.2018.04.015

19. Bornstein C, Nevo S, Giladi A, et al. Single-cell mapping of the thymic stroma identifies IL-25-producing tuft epithelial cells. Nature. Jul 2018;559(7715):622–626. doi:10.1038/s41586-018-0346-1

20. Miller CN, Proekt I, von Moltke J, et al. Thymic tuft cells promote an IL-4-enriched medulla and shape thymocyte development. Nature. Jul 2018;559(7715):627–631. doi:10.1038/s41586-018-0345-2

21. Park JE, Botting RA, Dominguez Conde C, et al. A cell atlas of human thymic development defines T cell repertoire formation. Science. Feb 21 2020;367(6480)doi:10.1126/science.aay3224

22. Bautista JL, Cramer NT, Miller CN, et al. Single-cell transcriptional profiling of human thymic stroma uncovers novel cellular heterogeneity in the thymic medulla. Nat Commun. Feb 17 2021;12(1):1096. doi:10.1038/s41467-021-21346-6

23. Zeng Y, Liu C, Gong Y, et al. Single-Cell RNA Sequencing Resolves Spatiotemporal Development of Pre-thymic Lymphoid Progenitors and Thymus Organogenesis in Human Embryos. Immunity. Nov 19 2019;51(5):930–948 e6. doi:10.1016/j.immuni.2019.09.008

24. Handel AE, Cheuk S, Dhalla F, et al. Developmental dynamics of the neural crest-mesenchymal axis in creating the thymic microenvironment. Sci Adv. May 13 2022;8(19):eabm9844. doi:10.1126/sciadv.abm9844

25. Baran-Gale J, Morgan MD, Maio S, et al. Ageing compromises mouse thymus function and remodels epithelial cell differentiation. Elife. Aug 25 2020;9doi:10.7554/eLife.56221

26. Aad G, Abajyan T, Abbott B, et al. Measurement of top quark polarization in top-antitop events from proton-proton collisions at radicals=7 TeV using the ATLAS detector. Phys Rev Lett. Dec 6 2013;111(23):232002. doi:10.1103/PhysRevLett.111.232002

27. Ragazzini R, Boeing S, Zanieri L, et al. Defining the identity and the niches of epithelial stem cells with highly pleiotropic multilineage potency in the human thymus. Dev Cell. Nov 20 2023;58(22):2428–2446 e9. doi:10.1016/j.devcel.2023.08.017

28. Farley AM, Morris LX, Vroegindeweij E, et al. Dynamics of thymus organogenesis and colonization in early human development. Development. May 2013;140(9):2015–26. doi:10.1242/dev.087320

29. Corbeaux T, Hess I, Swann JB, Kanzler B, Haas-Assenbaum A, Boehm T. Thymopoiesis in mice depends on a Foxn1-positive thymic epithelial cell lineage. Proc Natl Acad Sci U S A. Sep 21 2010;107(38):16613–8. doi:10.1073/pnas.1004623107

30. Flanagan SP. ’Nude’, a new hairless gene with pleiotropic effects in the mouse. Genet Res. Dec 1966;8(3):295–309. doi:10.1017/s0016672300010168

31. Nowell CS, Bredenkamp N, Tetelin S, et al. Foxn1 regulates lineage progression in cortical and medullary thymic epithelial cells but is dispensable for medullary sublineage divergence. PLoS Genet. Nov 2011;7(11):e1002348. doi:10.1371/journal.pgen.1002348

32. Munoz JJ, Tobajas E, Juara S, Montero S, Zapata AG. FoxN1 mediates thymic cortex-medulla differentiation through modifying a developmental pattern based on epithelial tubulogenesis. Histochem Cell Biol. Dec 2019;152(6):397–413. doi:10.1007/s00418-019-01818-z

33. Dooley J, Erickson M, Farr AG. An organized medullary epithelial structure in the normal thymus expresses molecules of respiratory epithelium and resembles the epithelial thymic rudiment of nude mice. J Immunol. Oct 1 2005;175(7):4331–7. doi:10.4049/jimmunol.175.7.4331

34. Dooley J, Erickson M, Roelink H, Farr AG. Nude thymic rudiment lacking functional foxn1 resembles respiratory epithelium. Dev Dyn. Aug 2005;233(4):1605–12. doi:10.1002/dvdy.20495

35. Cordier AC. Ultrastructure of the cilia of thymic cysts in “nude” mice. Anat Rec. Feb 1975;181(2):227–49. doi:10.1002/ar.1091810206

36. Shultz LD, Lyons BL, Burzenski LM, et al. Human lymphoid and myeloid cell development in NOD/LtSz-scid IL2R gamma null mice engrafted with mobilized human hemopoietic stem cells. J Immunol. May 15 2005;174(10):6477–89. doi:10.4049/jimmunol.174.10.6477

37. Zeleniak A, Wiegand C, Liu W, et al. De novo construction of T cell compartment in humanized mice engrafted with iPSC-derived thymus organoids. Nature methods. Oct 2022;19(10):1306–1319. doi:10.1038/s41592-022-01583-3

38. Miller AJ, Yu Q, Czerwinski M, et al. In Vitro and In Vivo Development of the Human Airway at Single-Cell Resolution. Dev Cell. Sep 28 2020;54(6):818. doi:10.1016/j.devcel.2020.09.012

39. Travaglini KJ, Nabhan AN, Penland L, et al. A molecular cell atlas of the human lung from single-cell RNA sequencing. Nature. Nov 2020;587(7835):619–625. doi:10.1038/s41586-020-2922-4

40. Busslinger GA, Weusten BLA, Bogte A, Begthel H, Brosens LAA, Clevers H. Human gastrointestinal epithelia of the esophagus, stomach, and duodenum resolved at single-cell resolution. Cell Rep. Mar 9 2021;34(10):108819. doi:10.1016/j.celrep.2021.108819

41. Metzger TC, Khan IS, Gardner JM, et al. Lineage tracing and cell ablation identify a post-Aire-expressing thymic epithelial cell population. Cell Rep. Oct 17 2013;5(1):166–79. doi:10.1016/j.celrep.2013.08.038

42. Michelson DA, Hase K, Kaisho T, Benoist C, Mathis D. Thymic epithelial cells co-opt lineage-defining transcription factors to eliminate autoreactive T cells. Cell. Jul 7 2022;185(14):2542–2558 e18. doi:10.1016/j.cell.2022.05.018

43. Pestka S, Krause CD, Walter MR. Interferons, interferon-like cytokines, and their receptors. Immunol Rev. Dec 2004;202:8–32. doi:10.1111/j.0105-2896.2004.00204.x

44. Der SD, Zhou A, Williams BR, Silverman RH. Identification of genes differentially regulated by interferon alpha, beta, or gamma using oligonucleotide arrays. Proc Natl Acad Sci U S A. Dec 22 1998;95(26):15623–8. doi:10.1073/pnas.95.26.15623

45. Darnell JE, Jr., Kerr IM, Stark GR. Jak-STAT pathways and transcriptional activation in response to IFNs and other extracellular signaling proteins. Science. Jun 3 1994;264(5164):1415–21. doi:10.1126/science.8197455

46. Pfeffer LM. The role of nuclear factor kappaB in the interferon response. J Interferon Cytokine Res. Jul 2011;31(7):553–9. doi:10.1089/jir.2011.0028

47. Aibar S, Gonzalez-Blas CB, Moerman T, et al. SCENIC: single-cell regulatory network inference and clustering. Nature methods. Nov 2017;14(11):1083–1086. doi:10.1038/nmeth.4463

48. Van de Sande B, Flerin C, Davie K, et al. A scalable SCENIC workflow for single-cell gene regulatory network analysis. Nat Protoc. Jul 2020;15(7):2247–2276. doi:10.1038/s41596-020-0336-2

49. Matsumoto H, Scicluna BP, Jim KK, et al. HIVEP1 Is a Negative Regulator of NF-kappaB That Inhibits Systemic Inflammation in Sepsis. Frontiers in immunology. 2021;12:744358. doi:10.3389/fimmu.2021.744358

50. Cook ME, Jarjour NN, Lin CC, Edelson BT. Transcription Factor Bhlhe40 in Immunity and Autoimmunity. Trends Immunol. Nov 2020;41(11):1023–1036. doi:10.1016/j.it.2020.09.002

51. Seo H, Gonzalez-Avalos E, Zhang W, et al. BATF and IRF4 cooperate to counter exhaustion in tumor-infiltrating CAR T cells. Nat Immunol. Aug 2021;22(8):983–995. doi:10.1038/s41590-021-00964-8

52. Candi E, Rufini A, Terrinoni A, et al. DeltaNp63 regulates thymic development through enhanced expression of FgfR2 and Jag2. Proc Natl Acad Sci U S A. Jul 17 2007;104(29):11999–2004. doi:10.1073/pnas.0703458104

53. Silva FN, Albeshri A, Thayananthan V, Alhalabi W, Fortunato S. Robustness modularity in complex networks. Phys Rev E. May 2022;105(5-1):054308. doi:10.1103/PhysRevE.105.054308

54. Tandon A, Albeshri A, Thayananthan V, Alhalabi W, Radicchi F, Fortunato S. Community detection in networks using graph embeddings. Phys Rev E. Feb 2021;103(2-1):022316. doi:10.1103/PhysRevE.103.022316

55. Traag VA, Waltman L, van Eck NJ. From Louvain to Leiden: guaranteeing well-connected communities. Sci Rep. Mar 26 2019;9(1):5233. doi:10.1038/s41598-019-41695-z

56. Lottini G, Baggiani M, Chesi G, et al. Zika virus induces FOXG1 nuclear displacement and downregulation in human neural progenitors. Stem Cell Reports. Jul 12 2022;17(7):1683–1698. doi:10.1016/j.stemcr.2022.05.008

57. Steimle V, Siegrist CA, Mottet A, Lisowska-Grospierre B, Mach B. Regulation of MHC class II expression by interferon-gamma mediated by the transactivator gene CIITA. Science. Jul 1 1994;265(5168):106–9. doi:10.1126/science.8016643

58. Platanias LC. Mechanisms of type-I- and type-II-interferon-mediated signalling. Nat Rev Immunol. May 2005;5(5):375–86. doi:10.1038/nri1604

59. Theofilopoulos AN, Baccala R, Beutler B, Kono DH. Type I interferons (alpha/beta) in immunity and autoimmunity. Annu Rev Immunol. 2005;23:307–36. doi:10.1146/annurev.immunol.23.021704.115843

60. Boehm U, Klamp T, Groot M, Howard JC. Cellular responses to interferon-gamma. Annu Rev Immunol. 1997;15:749–95. doi:10.1146/annurev.immunol.15.1.749

61. Cunningham TJ, Duester G. Mechanisms of retinoic acid signalling and its roles in organ and limb development. Nat Rev Mol Cell Biol. Feb 2015;16(2):110–23. doi:10.1038/nrm3932

62. Duester G. Retinoic acid synthesis and signaling during early organogenesis. Cell. Sep 19 2008;134(6):921–31. doi:10.1016/j.cell.2008.09.002

63. Kopinke D, Sasine J, Swift J, Stephens WZ, Piotrowski T. Retinoic acid is required for endodermal pouch morphogenesis and not for pharyngeal endoderm specification. Dev Dyn. Oct 2006;235(10):2695–709. doi:10.1002/dvdy.20905

64. Chelbi-Alix MK, Pelicano L. Retinoic acid and interferon signaling cross talk in normal and RA-resistant APL cells. Leukemia. Aug 1999;13(8):1167–74. doi:10.1038/sj.leu.2401469

65. Rhinn M, Dolle P. Retinoic acid signalling during development. Development. Mar 2012;139(5):843–58. doi:10.1242/dev.065938

66. Kam RK, Deng Y, Chen Y, Zhao H. Retinoic acid synthesis and functions in early embryonic development. Cell Biosci. Mar 22 2012;2(1):11. doi:10.1186/2045-3701-2-11

67. DaFonseca CJ, Shu F, Zhang JJ. Identification of two residues in MCM5 critical for the assembly of MCM complexes and Stat1-mediated transcription activation in response to IFN-gamma. Proc Natl Acad Sci U S A. Mar 13 2001;98(6):3034–9. doi:10.1073/pnas.061487598

68. Zhang JJ, Zhao Y, Chait BT, et al. Ser727-dependent recruitment of MCM5 by Stat1alpha in IFN-gamma-induced transcriptional activation. EMBO J. Dec 1 1998;17(23):6963–71. doi:10.1093/emboj/17.23.6963

69. Goldfarb Y, Kadouri N, Levi B, et al. HDAC3 Is a Master Regulator of mTEC Development. Cell Rep. Apr 19 2016;15(3):651–665. doi:10.1016/j.celrep.2016.03.048

70. Stritesky GL, Xing Y, Erickson JR, et al. Murine thymic selection quantified using a unique method to capture deleted T cells. Proc Natl Acad Sci U S A. Mar 19 2013;110(12):4679–84. doi:10.1073/pnas.1217532110

71. Klein L, Kyewski B, Allen PM, Hogquist KA. Positive and negative selection of the T cell repertoire: what thymocytes see (and don’t see). Nat Rev Immunol. Jun 2014;14(6):377–91. doi:10.1038/nri3667

72. Anderson MS, Venanzi ES, Klein L, et al. Projection of an immunological self shadow within the thymus by the aire protein. Science. Nov 15 2002;298(5597):1395–401. doi:10.1126/science.1075958

73. Dhalla F, Baran-Gale J, Maio S, Chappell L, Hollander GA, Ponting CP. Biologically indeterminate yet ordered promiscuous gene expression in single medullary thymic epithelial cells. EMBO J. Jan 2 2020;39(1):e101828. doi:10.15252/embj.2019101828

74. Dermadi D, Bscheider M, Bjegovic K, et al. Exploration of Cell Development Pathways through High-Dimensional Single Cell Analysis in Trajectory Space. iScience. Feb 21 2020;23(2):100842. doi:10.1016/j.isci.2020.100842

75. Street K, Risso D, Fletcher RB, et al. Slingshot: cell lineage and pseudotime inference for single-cell transcriptomics. BMC Genomics. Jun 19 2018;19(1):477. doi:10.1186/s12864-018-4772-0

76. Van den Berge K, Roux de Bezieux H, Street K, et al. Trajectory-based differential expression analysis for single-cell sequencing data. Nat Commun. Mar 5 2020;11(1):1201. doi:10.1038/s41467-020-14766-3

77. Akiyama T, Shimo Y, Yanai H, et al. The tumor necrosis factor family receptors RANK and CD40 cooperatively establish the thymic medullary microenvironment and self-tolerance. Immunity. Sep 19 2008;29(3):423–37. doi:10.1016/j.immuni.2008.06.015

78. Boehm T, Scheu S, Pfeffer K, Bleul CC. Thymic medullary epithelial cell differentiation, thymocyte emigration, and the control of autoimmunity require lympho-epithelial cross talk via LTbetaR. J Exp Med. Sep 1 2003;198(5):757–69. doi:10.1084/jem.20030794

79. Burkly L, Hession C, Ogata L, et al. Expression of relB is required for the development of thymic medulla and dendritic cells. Nature. Feb 9 1995;373(6514):531–6. doi:10.1038/373531a0

80. Kinoshita D, Hirota F, Kaisho T, et al. Essential role of IkappaB kinase alpha in thymic organogenesis required for the establishment of self-tolerance. J Immunol. Apr 1 2006;176(7):3995–4002. doi:10.4049/jimmunol.176.7.3995

81. Pelicano L, Li F, Schindler C, Chelbi-Alix MK. Retinoic acid enhances the expression of interferon-induced proteins: evidence for multiple mechanisms of action. Oncogene. Nov 6 1997;15(19):2349–59. doi:10.1038/sj.onc.1201410

82. Matikainen S, Ronni T, Hurme M, Pine R, Julkunen I. Retinoic acid activates interferon regulatory factor-1 gene expression in myeloid cells. Blood. Jul 1 1996;88(1):114–23.

83. Kolla V, Lindner DJ, Xiao W, Borden EC, Kalvakolanu DV. Modulation of interferon (IFN)-inducible gene expression by retinoic acid. Up-regulation of STAT1 protein in IFN-unresponsive cells. J Biol Chem. May 3 1996;271(18):10508–14. doi:10.1074/jbc.271.18.10508

84. Farr AG, Rudensky A. Medullary thymic epithelium: a mosaic of epithelial “self”? J Exp Med. Jul 6 1998;188(1):1–4. doi:10.1084/jem.188.1.1

85. Kaiser C, Bradu A, Gamble N, Caldwell JA, Koh AS. AIRE in context: Leveraging chromatin plasticity to trigger ectopic gene expression. Immunol Rev. Jan 2022;305(1):59–76. doi:10.1111/imr.13026

86. Takaba H, Morishita Y, Tomofuji Y, et al. Fezf2 Orchestrates a Thymic Program of Self-Antigen Expression for Immune Tolerance. Cell. Nov 5 2015;163(4):975–87. doi:10.1016/j.cell.2015.10.013

87. Qi Y, Zhang R, Lu Y, Zou X, Yang W. Aire and Fezf2, two regulators in medullary thymic epithelial cells, control autoimmune diseases by regulating TSAs: Partner or complementer? Frontiers in immunology. 2022;13:948259. doi:10.3389/fimmu.2022.948259

88. Rode I, Rodewald HR. Transcription factor hijacking in the name of tolerance. Cell. Jul 7 2022;185(14):2398–2400. doi:10.1016/j.cell.2022.06.026

89. Barnes JL, Yoshida M, He P, et al. Early human lung immune cell development and its role in epithelial cell fate. Sci Immunol. Dec 15 2023;8(90):eadf9988. doi:10.1126/sciimmunol.adf9988

90. Hern WM. Correlation of fetal age and measurements between 10 and 26 weeks of gestation. Obstet Gynecol. Jan 1984;63(1):26–32.

91. Hafemeister C, Satija R. Normalization and variance stabilization of single-cell RNA-seq data using regularized negative binomial regression. Genome Biol. Dec 23 2019;20(1):296. doi:10.1186/s13059-019-1874-1

92. Stuart T, Butler A, Hoffman P, et al. Comprehensive Integration of Single-Cell Data. Cell. Jun 13 2019;177(7):1888–1902 e21. doi:10.1016/j.cell.2019.05.031

93. Moerman T, Aibar Santos S, Bravo Gonzalez-Blas C, et al. GRNBoost2 and Arboreto: efficient and scalable inference of gene regulatory networks. Bioinformatics. Jun 1 2019;35(12):2159–2161. doi:10.1093/bioinformatics/bty916

